# Binding Mechanism of the Matrix Domain of HIV-1 Gag on Lipid Membranes

**DOI:** 10.1101/2020.05.06.080945

**Authors:** V. Monje-Galvan, Gregory A. Voth

## Abstract

Aggregation of the HIV-1 Gag protein onto the plasma membrane (PM) enables viral budding and infection propagation. Gag assembly at the membrane interface is mediated by its matrix domain (MA), the Myristoylated (Myr) N-terminus. MA targets the PM through electrostatic interactions, mainly at its highly-basic-region (HBR). The mechanism of Myr insertion and its role in protein-membrane dynamics remains unclear. Using all-atom molecular dynamics, we examined an MA unit in the vicinity of lipid bilayers that model different characteristics of the PM. Interaction with PIP_2_ and PS lipids is highly favored around the HBR, and is enough to keep the protein bound. Additionally, we simulated three MA units near our bilayers and quantified the collective effects of free monomers vs. formed trimers on Myr insertion events. Micro-second-long trajectories allowed us to observe Myr insertion, propose a mechanism, quantify specific interactions with lipids, and examine the response of the local membrane environment.

## I. Introduction

Protein-lipid interactions play key roles in cell growth, signaling processes, and disease onset and propagation (1, 2). Protein-protein interactions themselves are frequently dependent or mediated by specific lipids in the membrane (3, 4); at the same time they modify the local environment of the binding site to propagate a process or signaling cascade (5-7). Peripheral membrane proteins have received renewed attention over the past decade as they are involved in membrane deformation processes (8), lipid or small molecule transport between membranes (5, 9, 10), as well as viral assembly (6). Despite these advances, there are still unresolved questions in terms of the specificity of peripheral proteins to cell organelles, anionic lipids, and lipids rafts or domains (11-13).

In the context of retroviral assembly, key peripheral proteins assemble at the membrane interface of an infected host, or can even start assembly in the cytosol and then target the plasma membrane (PM) for subsequent release (14, 15). Protein multimerization at the PM leads to the formation of an immature lattice and lateral reorganization on the membrane that result sin the budding and release of an immature virion enclosing the viral genetic code (16). Given the relevance of protein interactions at the membrane interface for viral replication, it is important to understand the mechanisms of membrane targeting by proteins as well as the role of specific lipid species in protein binding and aggregation.

One of the most studied retroviruses is the human immunodeficiency virus type 1 (HIV-1). A key constituent of this virus is the <u>g</u>roup specific <u>a</u>nti<u>g</u>en (Gag) polyprotein, which targets the PM and initiates the viral assembly process upon oligomerization at this interface (15). Membrane targeting of Gag is mediated by its N-terminus, the matrix (MA) domain (17) – see Fig. The lipidated tail of MA, a myristoyl (Myr), is conserved across most retroviruses and is proposed to be key for interactions with lipids in the PM (18). Several studies have examined and provided key conclusions in terms of protein-protein interactions to characterize the membrane targeting, viral assembly, and budding processes (16, 19-22).

Among the main challenges in the study of membrane targeting domains, like MA, is the transient nature of their interaction and conformational changes at the membrane interface (23). Advances in experimental and computational techniques have enabled the study of such interactions in the context of viral replication (3, 17, 18, 24). In recent years, the specificity of the MA domain to the inner leaflet of the PM has been studied, showing that MA binds the PM through electrostatic and hydrophobic interactions mainly from its highly-basic region (HBR), located near the Myr site (18). Different factors in membrane character and composition have been identified to enhance or reduce MA binding affinity to the bilayer (3). It has been shown that MA-PM interactions are far more complex than mere lipid headgroup recognition, and that protein binding also modifies membrane dynamics (15). For example, MA aggregation is needed for the recruitment of lipids and other viral proteins to the viral assembly site in HIV-1 (13, 25, 26). Additionally, the role of lipid rafts in protein-membrane interactions has been explored (27). Nonetheless, the specific mechanism and conformational changes of MA that enable protein binding and aggregation remain unclear. The mechanism of Myr exposure and insertion as it pertains to dynamics of the viral assembly site at the PM is also not yet well understood. Finally, the effect of MA binding on lipid reorganization at the PM and how that renders the binding site more suitable for the recruitment of additional protein units of the viral bud has not been characterized in detail (15, 28).

Here we summarize an all-atom molecular dynamics (MD) study designed to gain molecular insight into the MA-membrane interactions. It is established that lipid composition determines the character of a bilayer, and in turn, protein interaction and function (29); thus, studying processes at the membrane interface requires appropriate models to mimic the membrane environment, surface charge, and topology. Hence, we use three symmetric membrane bilayers to model different aspects of the PM mimicking the composition of experimental studies of HIV-1 Gag (20). Our models contain between 3 and 6 different lipid types at relevant ratios to model the PM. The following sections report our methods and relevant conclusions for MA binding and Myr insertion mechanisms.

## II. Results

### MA bound conformations

Two bound conformations, mouth-down (*blocked*) and mouth-to-the-side (*open*), occur within the first 100ns of the short equilibration runs that started with MA at least 1 nm away from the membrane (see Fig. 2). The protein was positioned with its mouth pointing downwards and Myr sequestered in the hydrophobic cavity of the protein; Table S1 summarizes the details of the systems presented in this study. The protein was free to move in the solvent and the final bound conformation occurs indistinctly to the top or bottom leaflets of our symmetric models. The initial conformation of the protein did not influence the final bound state. Table S2 summarizes the time of first contact with the bilayer as well as that of Myr exposure from the systems bound in the open conformation. Initial contact times with the *raft* model are approximately half than with the *chol-free* or the *inner* models, and both bound conformations occur in the replicas run.

**Figure 1.**
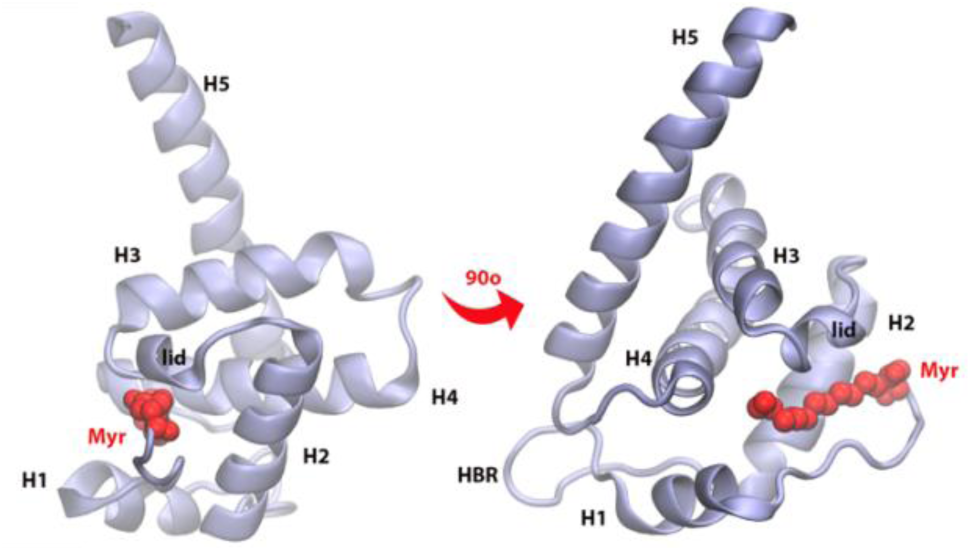
Matrix (MA) domain of HIV-1 Gag (residues 1-120) with key regions labeled: H1, H2, H3, H4, and H5 are helices numbered in order of appearance in the protein sequence; Myr is the lipidated tail of MA attached covalently to the N-terminus of the protein (shown in red); the lid is a small helical region that locks Myr inside the protein hydrophobic pocket; the HBR is a coiled region rich in LYS and ARG residues that is the main contributor to electrostatic interactions with the membrane.

**Figure 2.**
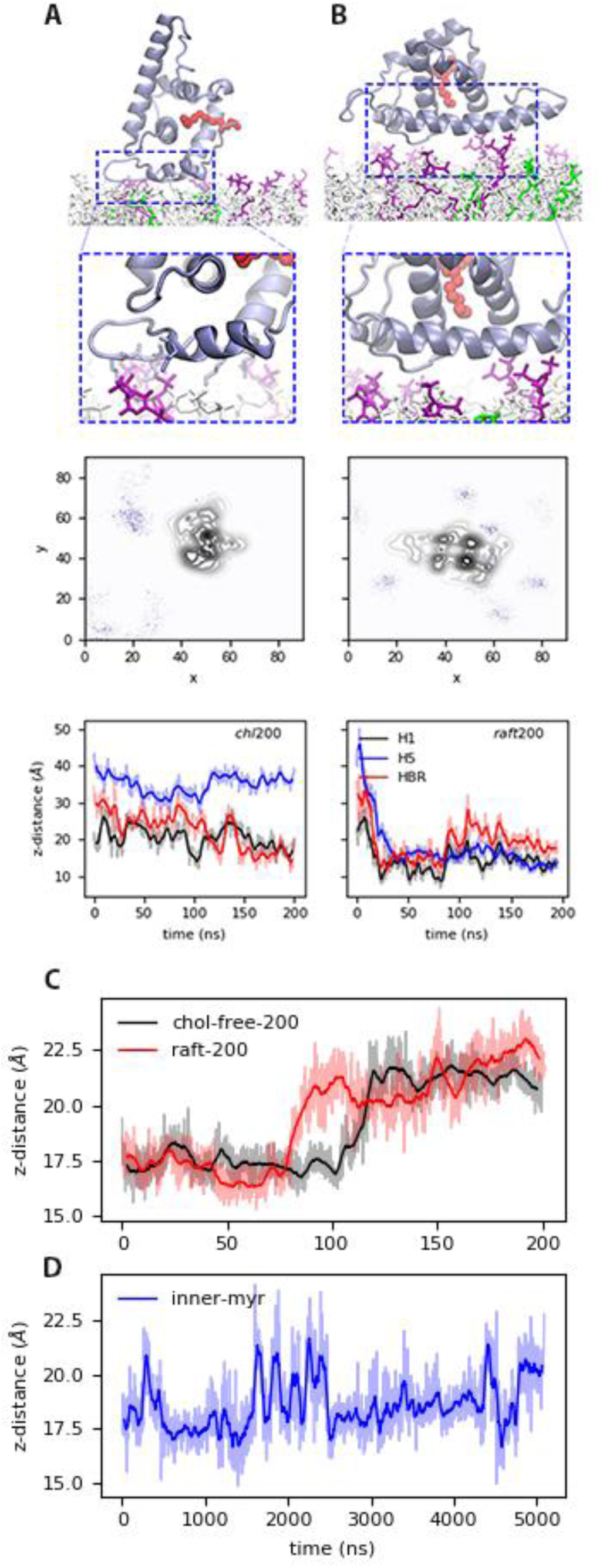
Bound states of MA with model membranes: (A) In the open state with the *Chol-free*_*200*_ model, and (B) the blocked state with the *Raft*_*200*_ model. <u>Top</u>: snapshots of the bound conformations, PIP_2_ lipids are shown in purple and DOPS in green. <u>Center</u>: PIP_2_ lipid density at the end of the 200ns trajectories. <u>Bottom</u>: distance between protein sections and the phosphate region of the lipids in the binding leaflet for each conformation. Time series of the distance between H1 and Lid of MA, i.e. the mouth of the protein for the (C) *chol-free*_*200*_ and *raft*_*200*_ systems with an open-bound and a blocked-bound MA, respectively. (D) H1-Lid distance for the *inner*_*myr*_ model trajectory. Moving average for these plots is shown in bold lines every 5ns for the 200ns trajectories, and every 50ns for the microsecond trajectory in bold lines; the faded curves are the original dataset.

In Fig. 2 we illustrate the open and blocked bound conformations with the *chol-free* and the *raft* models, respectively. The open conformation is characterized by the interaction of H1 and the HBR with the membrane surface. In this conformation, Myr is free to exit the protein’s hydrophobic cavity, and does so without much resistance; in fact, Myr is exposed and interacts with the lipid headgroups within our 200ns trajectories, though no insertion is observed from MA monomers until past the microsecond mark. Myr exposure occurs when the distance between the Lid and H1 regions, the mouth of the protein, opens up as shown in Fig 2.C for the *chol-free* and *raft* trajectories. On the other hand, the blocked conformation traps the Myr tail inside the protein hydrophobic cavity, which restrains its movement and prevents its insertion into the bilayer as it is discussed in subsequent sections. It is important to note the blocked conformation is not physically relevant when examining early stages of viral assembly, given the presence of the full Gag polyprotein would not allow for the interaction of H5 with the membrane as depicted in Fig 2.B. The blocked conformation could be a relevant structure of MA on the membrane surface during virus maturation, after budding and release of the virion and proteolytic cleavage of the MA-Capsid connection in the full Gag sequence (30).

In both conformations the electrostatic interactions are predominantly between LYS and ARG residues and DOPS or PIP_2_ lipids in the bilayer. The central panels in Fig. 2 show the lipid density of PIP_2_ at the end of the 200ns runs, averaged over the last 50ns of simulation. In the open conformation interactions with anionic lipids occur mainly with the HBR (Fig 2.A *chol-free* model), and with H5 in the blocked conformation (Fig 2.B, *raft* model). Fig S2. shows the lipid density of DOPS at the end of the corresponding trajectories; additionally, a sample with the *inner* model is shown in this figure for the trajectory with a protein bound in the open conformation. Both PIP_2_ and DOPS lipids co-localize to the protein binding site in the open conformation, but initially only PIP_2_ lipids interact with a protein bound in the blocked conformation. As will be discussed later, DOPS lipids interactions with the protein bound in the blocked conformation are slightly enhanced when Myr is inserted (see Fig S2 and S3.B). The protein is also able to sample both bound conformations when Myr is pre-inserted, which is partially modulated by contacts between the HBR and PIP_2_ lipids (see Fig S3.C).

### Myristate Insertion

We observed insertion of the Myr lipid tail repeatedly among our microsecond trajectories; Table 1 summarizes the systems in which this process is observed, and Table S3 lists the simulation length, protein bound conformation, and Myr location for all of these systems. Myr insertion occurred in two of the four trajectories of an MA monomer on the surface of a bilayer, simulated for 5 μ*s*; and in the simulation with a formed trimer in the case of multiple MA units on the membrane surface, simulated for 1 μ*s*. The systems with three separate MA monomers on the surface did not exhibit any Myr insertion within the 1 μ*s* trajectory. In all cases, insertion occurs only from the open bound conformation, i.e. the H1 and HBR interacting with the bilayer and an extended H5. Myr always unrolled into the binding leaflet and only occurred with the *inner* membrane model. Myr insertion never takes places from a blocked bound conformation, in which the lipid tail can roll on itself up and down the hydrophobic cavity of the protein, but cannot dive into the binding leaflet. Myr insertion is never observed with the *raft* models, not even from the open bound conformations of MA monomers on the surface of large membrane patches, or from a formed MA trimer.

**Table 1.** Summary of Myr insertion events for the systems that exhibit Myr insertion. The angle reported in the last column is that between the Myr tail and the bilayer normal, taken as the positive z-axis. The standard error is reported along with the angles.

| System | Sim time (us) | Bound conf. | Bound leaflet | insertion (ns) | Myr angle (°) |
| --- | --- | --- | --- | --- | --- |
| Inner | 5 | open | bottom | 1380 ± 2 | 25.4 ± 2.3 |
| Inner1 | 1.6 | open | bottom | 78 ± 1 | 8.2 ± 1.4 |
| Inner-*L<sub>trimer</sub>* | 1 | open | top | MA1: 350 ± 4<br>MA2: 780 ± 2<br>MA3: none | MA1: 37.8 ± 5.4<br>MA2: 18.1 ± 3.2<br>MA3: N/A |

Figure 3.A shows a series of snapshots showcasing the insertion of Myr into the *inner* membrane model, a process that occurs between 70-180ns when a single MA protein is present. The first carbon in the tail, C2, interacts closely with the lipid headgroups upon exposure from the hydrophobic cavity of the protein; the tail can lay flat on the membrane surface while it searches for a large-enough lipid packing defect suitable for insertion. During insertion, the last carbon of the tail, C14, can extend vertically towards the water; as the tail rolls into the binding leaflet, C14 gets close to C2 and then extends into the center of the bilayer. The mechanism is similar when a formed MA trimer is on the surface of a bilayer, but in this case Myr can also rest flat between the protein and the membrane, and unroll into the leaflet from this position. Again, Myr insertion never occurs in a diving fashion. Insertion was permanent on the timescale of the simulation in the case of MA monomers on the surface of small membrane patches; from an MA trimer, the first Myr insertion occurs before the 500ns mark, but the tail exited immediately and inserted permanently after 700ns.

**Figure 3.**
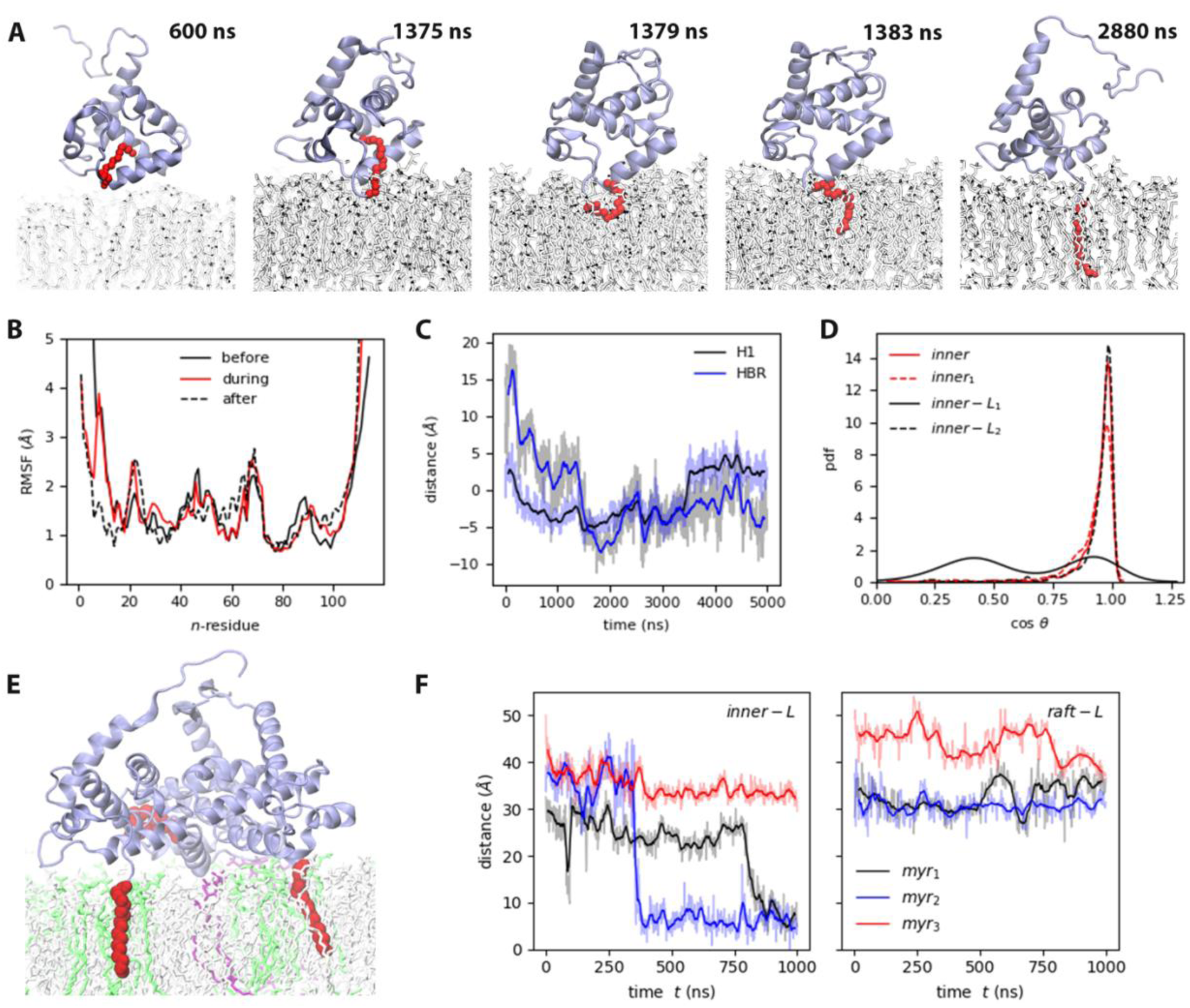
(A) Myristate insertion mechanism (from the *inner* trajectory). (B) RMSF of MA before, during, and after Myr insertion, at least 50ns of the respective sections in the *inner* trajectory were used for this analysis. (C) Self-distance of H1 and HBR during Myr insertion (refer to Table S4 for the definition of *self-distance*). (D) Probability distribution of the angle between Myr and the bilayer normal for the MA units that exhibit Myr insertion. (E) Snapshot of the side view of *inner-L*_*trimer*_ at the end of the simulation, PIP_2_ lipids are shown in purple, DOPS lipids in green. (F) Time series of Myr insertion in the first and second MA units in the *inner-L*_*trimer*_ simulation, the distance was computed between the last carbon in the lipid tail (C14) and the bilayer center. All the time series show the blocked moving averaged in solid color and the raw data faded.

Fig 3.B shows the root-mean-squared-fluctuations (RMSF) of the protein before, during and after Myr insertion for the *inner*-MA system. Note that residues 2-7, 19-29, 44-46, 52, 69-71, and 90-95 fluctuate the most, as is expected for coiled regions. However, residues 8-18 that correspond to H1, fluctuate more before and during Myr insertion, and stabilize afterwards. As Myr inserts, H1 gets closer to the bilayer and lays flat on the membrane surface; this can be observed in the time series inset on Fig 3.B that shows the self-distance of both H1 and HBR computed as the z-component of the distance between the first and last alpha-carbons of each region (refer to Table S4 for this metric). Upon insertion, Myr aligns to the lipid tails and remains in this position until the end of the simulation. Fig. 3.C shows the probability distribution of the cosine of the angle between Myr and the bilayer normal; in all cases, the lipid tail remains aligned to the z-axis inside the bilayer (cos (*θ*) =1), and C14 moves up to the middle region of the leaflet occasionally.

To further examine the effects of Myr on the binding leaflet, we simulated an MA monomer on the surface of the small *inner* and *raft* membrane models starting from a pre-inserted Myr configuration. Similarly, we ran three replicas of this configuration with the *chol-free* model to access the effect of cholesterol, i.e., the mechanical and structural properties of the membrane, on Myr retention and insertion. Fig. S4 shows the probability distribution of the cosine of the angle between Myr and the z-axis for these systems. Note that in the *raft* model, a membrane rich in cholesterol and sphingomyelin lipids, the angle distribution is much narrower than in the more fluid *chol-free* model. The trajectory with the *inner* model started from a partially inserted Myr, but contrary to our expectations, the Myr tail exited the membrane and returned to the hydrophobic cavity of the protein.

Figure 4 shows three of the simulations with the *inner* model, the first one in which Myr insertion occurs from an MA monomer, the second one that started with a partially started Myr, and the last one from one of the MA units that exhibited Myr insertion from the trimer configuration. Notice the C2 carbon in Myr, the one attached to the GLY residue in the N-terminus of MA, does not shift much during insertion because it is already quite close to the membrane upon protein binding. The relative contribution to Myr conformation in the system with respect to the C2 distance to the bilayer center has only one minimum. C14, on the other hand, can stay inside the protein cavity at approximately 4 nm from the bilayer center, rest between the protein and the membrane surface around 3.7 nm from the center, or remain inserted between 0-0.6 nm from the center. Interestingly, when Myr returns to the hydrophobic cavity in lays nearly horizontally, aligned to the membrane surface inside the protein with C2 and C14 at the same height above the bilayer center. This is also observed in the probability distribution of the cosine of the angle between Myr and the z-axis for this system, included in Fig. S4 for comparison with the data for the *raft* and *chol-free* simulations that started with a pre-inserted tail.

**Figure 4.**
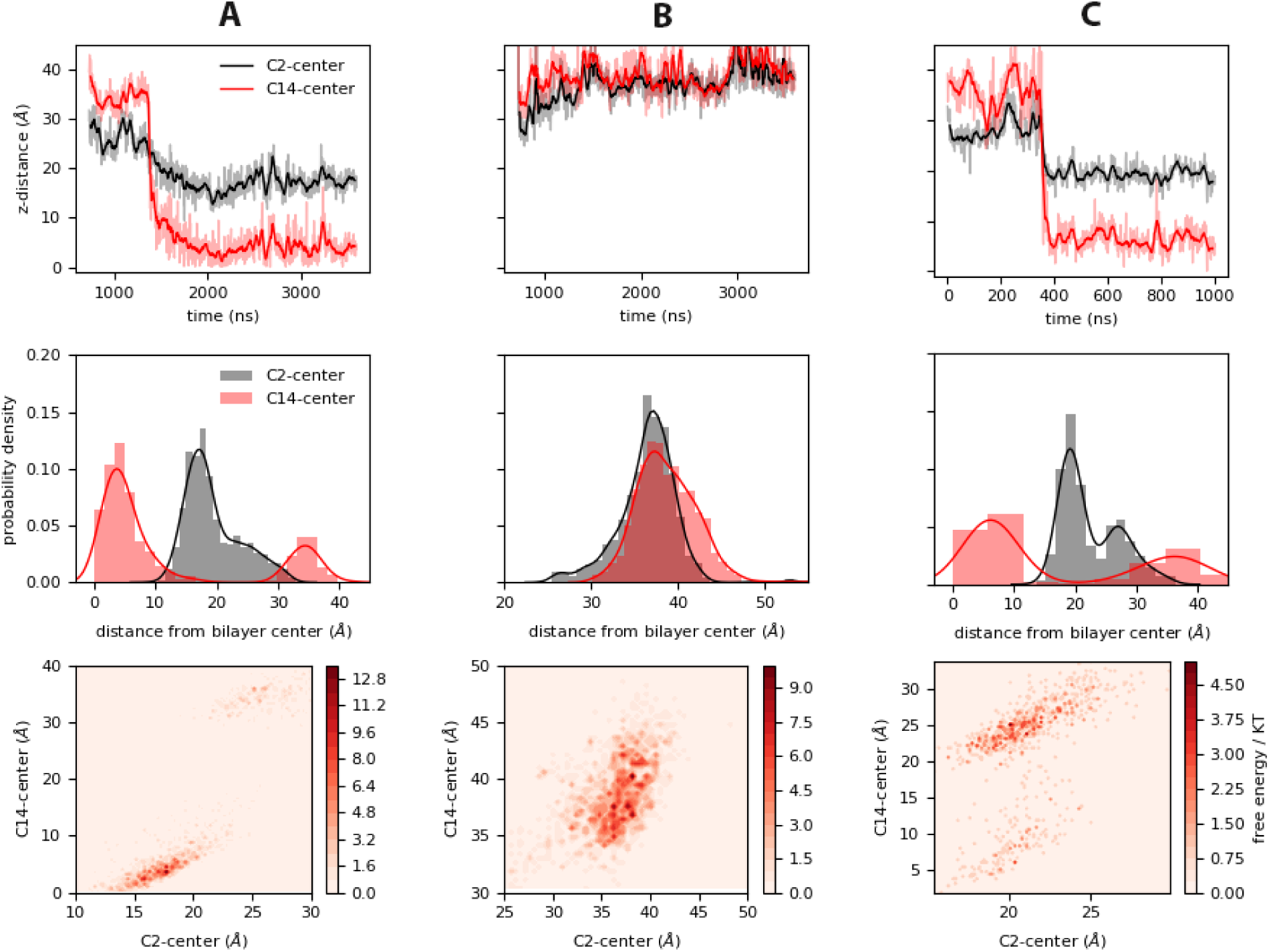
<u>Top</u>: Time series of insertion for the first and last carbons in the Myr tail, C2 and C14; the distance was computed between the center of mass of the respective carbon and the bilayer center as estimated from the average distance between the phosphate regions of the lipids in each leaflet. The faded regions show the raw data of this metric, and the bolded regions show the corresponding blocked moving average. <u>Center</u>: Probability density of the same distances as estimated from the normalized histograms of each time series. <u>Bottom</u>: Heat maps of the system with respect to the position of the C2 and C14 atoms of Myr as estimated from the histogrammed data. The systems shown are: (A) *Inner*, (B) *Inner*_*myr*_, and (C) *Inner-L*_*trimer*_ (MA_2_ unit).

### Membrane Response

As mentioned before, charged lipids co-localize to the binding site of the protein, even when only a single unit is present (refer to Fig. 2 and Fig. S2). PIP_2_ recruitment is not required for Myr insertion, but in some instances the number of contacts with MA increase upon Myr insertion as shown in Fig. 5 for the *inner* trajectories, and the *raft* model in Fig S3. In other cases, the number of contacts between MA and PIP_2_ remains the same or increases even without insertion of the lipid tail, as for the trajectories with the *raft* model in which the protein was bound in the blocked conformation (see Fig. S3). Note that MA can move more freely on the membrane surface after its initial binding and prior to Myr insertion, but is fixed in a given local lipid environment when starting from a pre-inserted Myr configuration. For the systems starting with a pre-inserted Myr, a maximum of 57% or 44% of PIP_2_ present in the *Chol-free* and *Raft* models, respectively colocalizes at the protein binding site. Whereas 75-100% of PIP_2_ lipids bind to the protein in the *Inner* model trajectories.

**Figure 5.**
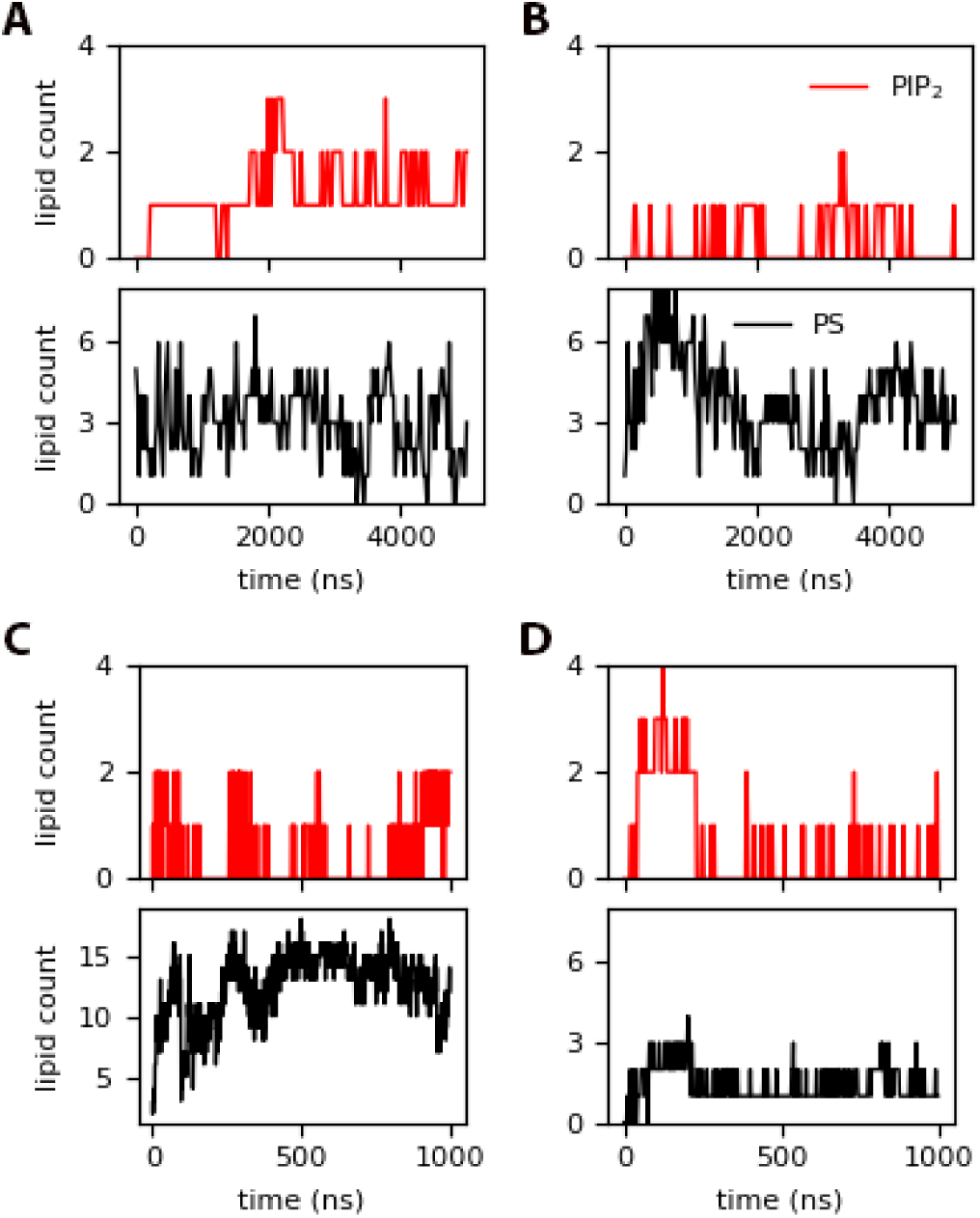
PIP_2_ and DOPS contacts within 10 Å of the protein for: (A) *inner* (insertion around 1380ns), (B) *inner*_*myr*_, (C) *inner-L*_*trimer*_ (with permanent Myr insertion after 400 ns), and (D) *raft-L*_*trimer*_ (with no insertion).

In addition to PIP_2_, DOPS lipids also interact with the protein; these are present in higher concentration in the *chol-free* and *inner* models, but at the same concentration as PIP_2_ in the *raft* model. When Myr is pre-inserted, the interaction of PS lipids with the protein increases from 12 to up to 30% of its content in the *raft* model, but fluctuates greatly between 9-35% of PS present in the *chol-free* model. From the trajectories with the *inner* model, PS-protein interactions are not affected by Myr insertion, and a maximum of 23% of PS present in the membrane interacts with the protein throughout the simulation. On the other hand, for the simulation that started with a partially inserted Myr, a maximum of 38% of PS present in the *inner* model interacts with the protein, and this decreases to 19% after Myr returns to the protein’s hydrophobic cavity.

In the case of formed MA trimers, as show in the bottom panels of Fig. 5, protein-DOPS interactions are enhanced after Myr insertion in the *Inner-L*_*trimer*_ model, but interactions with PIP_2_ do not follow a steady pattern. The trajectory with the *Raft-L*_*trimer*_ model shows that interactions with anionic lipids are not enhanced when Myr does not insert into the bilayer. Electrostatic interactions are still present between the protein and the membrane and are strong enough to keep the protein bound, but contacts with anionic lipids are reduced to nearly 50% when insertion does not occur. Additionally, Fig S5 shows lipid density plots for PIP_2_ and the relative locations of either the MA monomers or the formed MA trimer on the surface of the large membrane models. PIP_2_ co-localizes to the protein binding site in the same fashion as observed in the simulations of a single MA unit on the small membrane patches. As the monomers get closer together, PIP_2_ density further increases locally. The formed trimer simulated near the inner or raft models never dissociates from the bilayer, with or without Myr insertion.

In all the trajectories run for this study, Myr insertion was never observed into the leaflets of the *raft* model. However, we compared the effect of pre-inserted Myr in the aggregation of sphingomyelin lipids in the binding vs the opposite leaflet in the small membrane patches as well as the aggregation pattern of sphingomyelins in the *raft-L* membrane patch simulated with a formed MA trimer. We observe cholesterol aggregates at the same location as sphingomyelins as expected for these raft-related species. It is interesting to note sphingolipids do not co-localize to the protein binding site in the binding leaflet, but they seem to locate below this site in the opposite leaflet.

As a final metric, we computed the deuterium order parameters (**S**_**CD**_), a measure of order in terms of the alignment of lipid tails inside the bilayer, of the systems with Myr insertion events to membrane-only simulations of the *inner* membrane model (equilibrated on GROMACS for 750 ns). The S_CD_ values were computed per leaflet after Myr insertion for the protein-membrane systems and over the last 200ns of simulation for the membrane-only systems. In the protein-membrane systems the leaflets are identified as the binding (BL) and the opposite leaflets (OL) since the systems in this study are symmetric. The difference between the S_CD_ value of the protein-membrane and membrane-only systems is shown in Fig. 6. A positive number indicates the protein-membrane system has higher S_CD_ values, i.e. the lipids tails are more ordered inside the bilayer, while a negative value indicates more disorder in the lipid tails of the systems simulated with the protein vs the membrane only systems. The last carbons of the long tails of both PIP_2_ and sphingomyelin lipids, with 20 and 22 carbons respectively, experience an increase in order when the protein is present and Myr inserts into the bilayers. Moreover, the trend is different when a trimer is present instead of an MA monomer. This is particularly striking for the sphingomyelin values, which show an increase in order in the middle section of the lipid tail in the binding leaflet when a trimer is present instead of a monomer. On the other hand, the sphingomyelin lipids in the opposite leaflet show an even larger effect of disorder in the tail region, though the mid-section does experience an increase in order as in the binding leaflet larger than the case with a monomer on the membrane surface.

**Figure 6.**
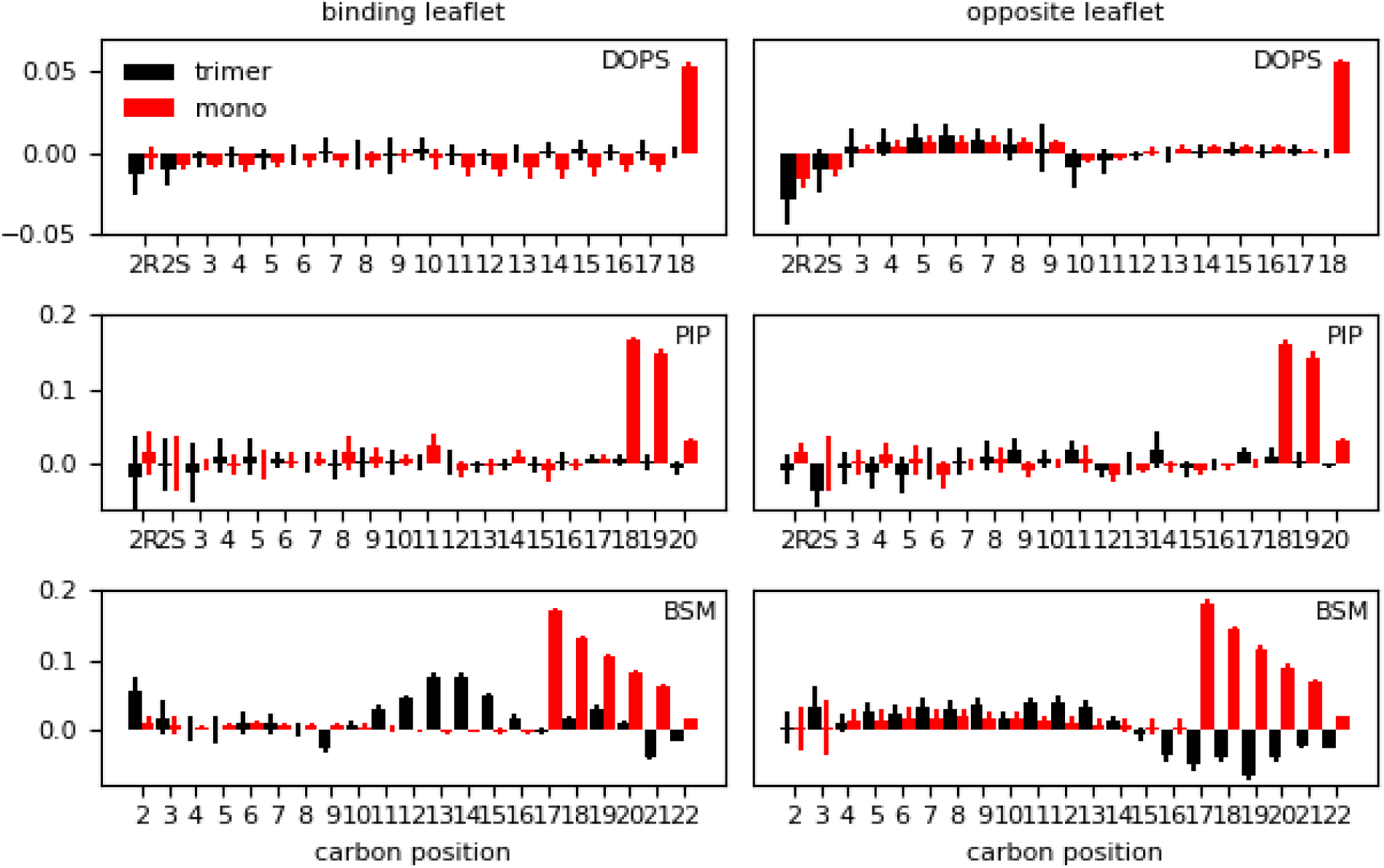
Difference in lipid order parameter (S_CD_) between the protein-membrane and membrane-only systems per leaflet for the *inner* (‘mono’) and *inner-L*_*trimer*_ (‘trimer’) systems after Myr insertion. A positive value indicates the presence of the protein with inserted Myr increases the order in the bilayer; whereas a negative value corresponds to a decrease in order with respect to the membrane-only system.

## III. Further Analysis and Discussion

We studied the effect of membrane composition on the binding mechanism and insertion of the lipidated tail of MA, the membrane targeting domain of HIV-1 Gag polyprotein. Using MD, we simulated MA monomers as well as formed trimer near the surface of different membrane models. These models were selected based on experimental studies that also examined the interplay between protein binding and aggregation on membranes in the context of viral assembly (20). Our models mimic the sterol content, surface charge, and key lipid species in the PM. Though we used a symmetric bilayer in this study, our models provide important insight on the protein-membrane interactions needed during the early stages of viral assembly. The key contributions from our studies include specific protein-lipid interactions at the binding site, a molecular-level mechanism for Myr insertion, characterization of protein conformational changes that enable Myr insertion, and the local changes in the membrane as a result of binding and insertion.

In agreement with experimental studies, the composition of the membrane affects and modulates the interaction of MA with the bilayer (3, 17, 31, 32). All of our models have a minimum of 12% anionic character, which is enough to promote electrostatic interactions with the HBR or H5 regions of the protein depending on bound conformation. Most of these interactions occur between LYS or ARG residues and PIP_2_ and DOPS lipids, as has also been stated in the literature (17, 23, 32). Figures 2, S2, & S3 show enrichment of PIP_2_ to the protein binding site as it has been suggested several times before (3, 20, 22, 33). Upon initial contact and binding, the protein is relatively free to search the membrane surface, as we observed on our microsecond trajectories with the *inner* and *raft* models with both MA monomers and a formed trimer. Insertion of the lipidated tail restricts the movement of MA, though it does not serve as the only anchor because the protein remains bound for the entirety of 3 or 5 μ*s* trajectories even in the cases where Myr insertion is not observed (*raft* model). Myr has been suggested as an anchor before, but experimental studies with Myr-less MA mutants also observed MA binding to membranes (3, 23). So far, there has not been a unified answer as to what is the role of Myr in MA membrane targeting and subsequent oligomerization of Gag at the inner leaflet of the PM. Below we propose specific MA-membrane interactions that enable exposure and insertion of Myr dependent on the membrane nature, which has not been captured through simulations previously.

We observe quick MA binding to our symmetric models in two conformations during the short simulation trajectories, shown in Fig 2 and summarized on Table S2. Both conformations are stabilized mainly by electrostatic interactions with PIP_2_ lipids, in agreement with various reports (17, 33). The protein orientation in the open conformation agrees with experimental and other simulation studies (23, 33). In this conformation H1 and HBR interact closely with the bilayer and H5 is extended towards the solvent; this is the only conformation from the two observed in our simulations that is plausible during early stages of viral assembly when the full Gag protein is present and connected to MA after H5. This conformation allows for intermittent interaction of H2 with the bilayer and seems to be the preferred orientation of MA. The open conformation is strongly modulated by HBR-PIP_2_ interactions, as shown in Fig S3.C for one trajectory with the *raft*_*pre*_ model. In this example, Myr started from a pre-inserted configuration and the protein bound in the blocked conformation, i.e., H5 flat on the membrane surface and HBR pointing towards the solvent. PIP_2_-mediated interactions with the HBR and a shift of H1, attached to Myr, into the lipid headgroup region enable the permanent transition from the blocked to the open conformation within the first microsecond of simulation until the end of the 5 μ*s* trajectory. It appears that the open bound conformation is the preferred orientation of MA, even without the presence of the full Gag in our studies.

Myr exposure is readily observed from the open conformation within the 200ns equilibration trajectories. The lipid tail exits the protein’s hydrophobic cavity as the Lid region separates from H1, opening the mouth of the protein. Figure 2.C&D shows time series of this distance corresponding to the two systems shown in Fig 1. Opening of the Lid-H1 region occurs in both the open and blocked bound conformations of MA, but only the open conformation allows for Myr exposure. In the blocked conformation, opening of the mouth only results in more room for Myr to wiggle inside the hydrophobic cavity of the protein; this configuration would allow for Myr diving into the membrane, but this mechanism is not observed in any of our trajectories – the preferred insertion mechanism is discussed in the following paragraphs. Opening of the Lid-H1 region occurs both for exposure and sequestration of Myr; in the trajectory of MA with the *inner* model starting from a partially inserted Myr, we see the lipid tail return to the hydrophobic cavity (Fig 4.B) as the mouth opens and closes to lock the tail back inside the protein (Fig 2.D).

Previous studies examined the protein in solution and suggested MA trimerization was needed for Myr exposure (34). Charlier *et al* (33) used umbrella sampling to identify a barrier close to 8 Kcal/mol for full Myr exposure in solvent. We conclude that the character of the membrane has a greater influence on protein-lipid interactions that allow Myr exposure and later insertion. In the short equilibrations we performed for a single MA or the MA trimer in solvent prior to positioning them above equilibrated membrane coordinates, we actually observe Myr sequestration into the hydrophobic cavity of the protein; which is expected intuitively as the hydrophobic tail would prefer a hydrophobic environment over surrounding water. Interestingly, one of our short protein-only equilibration systems did result on Myr leaning in the outer hydrophobic cavity of the protein, formed on the outer surface of the protein between the Lid and H3. We did not use this structure in any of our protein-membrane simulations, but it would be an interesting test case to examine the protein motions that enable ‘release’ of Myr from this external hydrophobic pocket. This external hydrophobic cavity is different from the location suggested from NMR studies to be a lodging pocket for membrane lipid tails to anchor MA to the membrane (18, 32), but suggests a lipid tail could potentially find an energetically favorable location outside the protein.

In itself, Myr exposure is not a determining step for insertion into the bilayer or protein retention on the membrane surface. We observe Myr insertion into our *inner* membrane model repeatedly in an unrolling fashion from an MA monomer or a formed trimer. In the trimer configuration, each protein unit is bound to the membrane in the open conformation, the mouth of the protein open side-wise and nearly perpendicular to the bilayer. We simulated trimers on the surface of the *inner* and *raft* models; nonetheless, no insertion is observed onto the *raft* model. Examining the nature of the membrane models, we see lipid packing in the bilayer as measured by the surface are per lipid (APL) is not a key determinant for Myr insertion. As listed on Table S1, the average APL is not statistically different between the *inner* and *raft* models. Key differences between these membrane models are the surface charge, 19% on the *inner* model vs 12% on the *raft* model, sphingomyelin content, 8% vs 30% respectively, and POPE content, 25% vs none respectively. We did not quantify lipid packing defects on our systems, but by simple inspection of the lipid species in each model, the *raft* model is expected to have more tightly packed lipid headgroups on its surface given the relative size of the headgroups in the lipid species in this model vs those in the *inner* model (see Fig. S1). Lastly, in the actual biological system, MA is not expected to bind a leaflet rich in cholesterol/sphingomyelin lipids, as is the outer leaflet of the PM. The fact Myr insertion is not observed onto the *raft* model also demonstrates the need for careful selection of lipids in the membrane models used to study protein interactions at the membrane surface, or inside the bilayer for that matter (29).

Considering the effect of Myr on lipid-lipid interactions, we observe sphingomyelin aggregates are more defined when Myr is inserted, for example, in the binding leaflet of the *Raft*_*pre*_ system. Furthermore, cholesterol molecules co-localize to the edge of sphingomyelin regions in the binding leaflet, but not inside them. Interestingly, these clusters are not below the protein binding site. The effect on lipid clustering is not as marked in the *inner* model runs with a single MA monomer given the sphingomyelin content is much lower than that of the *raft* model. Nonetheless, as shown in Fig S6, there are some sphingomyelin-cholesterol clusters in the *inner-L*_*trimer*_ trajectory. The presence of the trimer induces the formation of smaller or compact clusters that are again outside of the protein binding site. In this case however, cholesterol molecules can aggregate below the protein itself. An interesting note is that Myr seems to avoid fully saturated regions for its initial insertion; Figure 3.F shows the MA_1_ unit in the trimer partially inserted into the bilayer and exit immediately to reinsert permanently later in the trajectory. The initial insertion site was rich in cholesterol molecules, which made the lipid tail bounce back into the lipid headgroup region. Permanent insertion occurred when less cholesterol molecules were present, yet sterol molecules later accumulated around Myr afterwards. The dynamics of Myr sensing for permanent insertion remain to be fully elucidated; our results, however, suggest a feedback loop between Myr insertion events and local lipid composition.

As expected, no clustering is observed in the *chol-less* model because it lacks the lipids involved in domain/raft formation. Furthermore, these trajectories show no statistical difference in the lipid order parameters (S_CD_) compared to the corresponding membrane-only system even starting from a pre-inserted Myr configuration. Whereas these values in the *inner* membrane show an increase in order when the protein is present and Myr is inserted in both the monomer and trimer trajectories (see Fig. 6). Clearly, the membrane composition determines the extend of influence of the protein on lipid-lipid interactions. Capturing the changes in lipid dynamics on the membrane indicates Myr plays a role in lipid sorting and dynamics at the viral assembly site, rather that influencing the MA binding events. We did not quantify lipid clustering or interleaflet dynamics in this study, but have reserved this analysis for a future publication examining MA binding and aggregation on asymmetric bilayers.

As a final assessment, we characterized Myr insertion events using a reduction technique to quantify the slowest motions of the process. We performed tICA on the distances between each carbon of the Myr tail and the alpha-carbons of the structured regions of the protein as labeled on Fig 1: H1-5 and the HBR, no other coiled region was considered. Figure 7 shows two cases with the *inner* membrane model; panel A is for the trajectory exhibiting Myr insertion (*Inner* system) and panel B for the trajectory starting with a partially inserted Myr that resulted in its sequestration back into the protein’s hydrophobic cavity (*Inner*_*myr*_ system). Notice the first independent component (tIC) identified in the analysis seems to be rather related to protein motions for Myr exposure or sequestration, such as the opening and closing of the mouth of the protein, i.e., the H1-Lid distance (refer to Fig. 2.D). The second tIC captures Myr insertion, or lack thereof, depending on the trend of the time series. Fig. 7A shows a concave shape for the second tIC that decreases around the time of Myr insertion; whereas panel B shows a convex shape for the second tIC that increases as Myr is sequestered back into the protein around 2000ns, same time in which the H1-Lid region opens to make way for the lipid tail (see Fig. 2.C&D).

**Fig 7.**
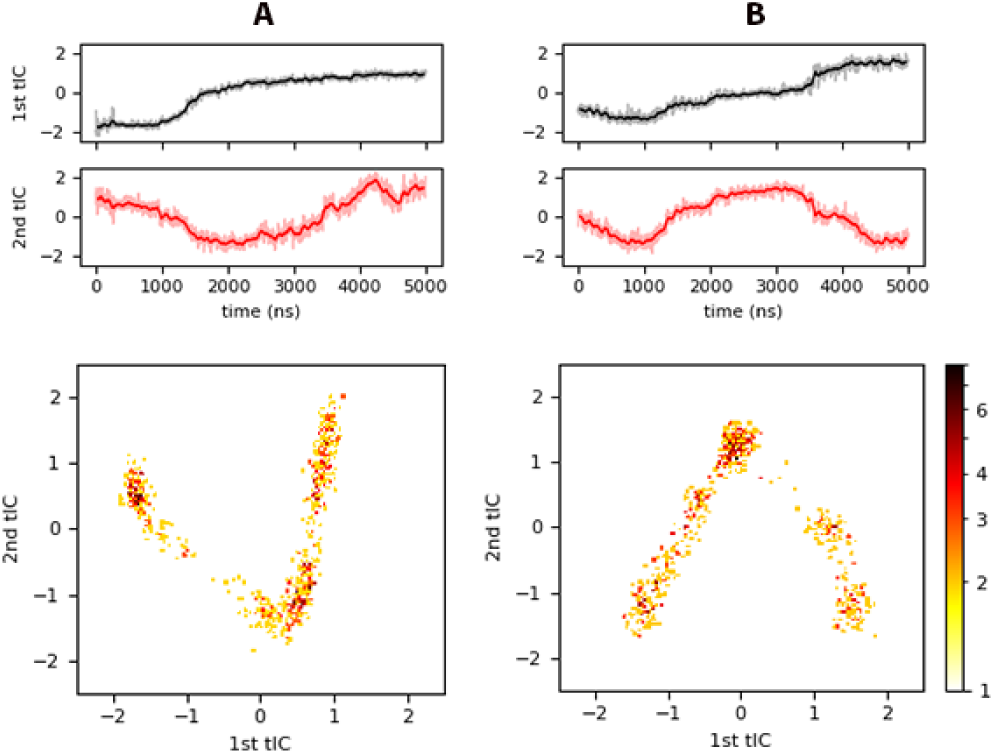
tICA results for (A) *inner*, and (B) inner_*myr*_. <u>Top</u> panels show the projection of each trajectory onto the slowest independent components identified, and the <u>bottom</u> panels a heat map of histograms of the projected trajectory onto the respective tIC (logarithmic scale).

The bottom panels in Fig 7 are 2D histograms of the trajectory projections onto the slowest tICs in logarithmic scale for clarity. These show a V-shape characterizes Myr insertion, while an inverted V characterizes the motion of Myr away from the bilayer center, either to the protein cavity or remaining around the lipid-headgroup region. The same characteristic shapes are observed for the corresponding plots of the *Chol-free*_*pre*_, *Raft*, and *Raft*_*pre*_ systems in Fig S7. In these cases, the systems with pre-inserted Myr (Fig S7.A & C) show a first tIC that increases in time, and a second tIC with a concave shape. Again, systems with an inserted Myr have a 2D histogram with V-shape behavior, while the system with no insertion shows an inverted V. Simply looking at the first tIC across the *inner, raft*, and *chol-free* models, it is apparent the intra-protein and protein-membrane interactions are different depending on the membrane model in use.

Examination of the loads of each tIC, i.e., the coefficients of each eigenvector, revealed the key distances that contribute to the slowest tIC are between the first two carbons in the Myr tail, C2 and C3, and the HBR, followed by the distances between the same carbons and H5; the distance between the middle carbon (C6) and H5 as well as the last carbon (C14) in the tail with respect to the HBR followed in third instance. In the case of the second slowest tIC, the most relevant distances were between the first carbons (C2 and C3) and the last ones (C13 and C14) of Myr and both the HBR and H1; additional contributions come from the C3-H5 and C13-H5 distances, and lastly from C6-H5 and C14-H4. These distances were deemed relevant because the respective normalized loads (coefficients) were higher than ± 0.5. The conclusions drawn from tICA also corroborate the preferred mechanism of Myr insertion by an unrolling of its tail, and provides further insight on key protein regions that contribute to the tail’s insertion.

## IV. Methods

Three membrane models were designed to study the binding dynamics of MA based on an experimental study from Yandrapali, *et al* (20). Table S1 in the Supporting Information (SI) summarizes the lipid components of each bilayer along with other system details. The *chol-free* model was selected to examine the impact of cholesterol on protein-membrane interactions. The *inner* model mimics the anionic character of the PM, while the *raft* model emphasizes the ability of the PM to phase separate and form lipid clusters enriched in sphingomyelin and cholesterol lipids.

Fully hydrated bilayers (35+ water molecules per lipid) and a MA protein were built separately using CHARMM-GUI *Membrane Builder* and *Quick-Solvator*, respectively (35-39). All systems were rendered electrostatically neutral using KCl salt at a 0.15 M concentration as summarized in Table S1. MD simulations were performed using the CHARMM36m force field (40), with its most up-to-date parameters for lipids and proteins. Initial equilibration of the bilayers and protein were performed using the GROMACS MD simulation package (41), and micro-second production runs were carried on the Anton 2 machine (42). Coordinates for an MA monomer were extracted from PDBID:1HIW in the protein data bank (https://www.rcsb.org) and a myristoyl (Myr) tail added to its N-terminus on CHARMM-GUI. Initial relaxation of the bilayer systems was carried out over 250 ps following the six-step protocol provided on CHARMM-GUI (43), and each system equilibrated for at least 100ns on GROMACS before introducing the protein in the solvent region. The protein, with an exposed Myr, was equilibrated separately for 150ns to allow the sequestration of Myr in the internal hydrophobic cavity of the protein. Equilibrated coordinates of each bilayer and the protein were merged with the protein at least 1.0 nm above the bilayer using GROMACS. The protein-membrane systems started with a sequestered Myr MA and were simulated for 200ns on GROMACS. Coordinates from equilibrated systems with the *inner* and *raft* models were extended to microsecond-trajectories on Anton2 to characterize Myr exposure and insertion events. Additionally, these bilayers were simulated starting from an inserted Myr configuration to examine its long-term effect on membrane dynamics. The *chol-free* model was also simulated on Anton2 starting from an inserted Myr configuration to compare the effects of the lipid tail on membrane properties when cholesterol is not present.

The equilibrated membranes were later replicated to form larger membrane patches to simulate three MA monomers or a formed MA trimer near the membrane surface. The smaller membrane patches contained 150 lipids per leaflet, while the larger patches have 600 lipids per leaflet with a surface of 282.24 nm_2_ (16.8 × 16.8 nm). The MA trimer (PDBID:1HIW) corresponding to the structure published by Hill *et al* (44) was equilibrated in water separately and then introduced near the membrane surface. During the trimer-only equilibration, two Myr tails in individual units were sequestered into the hydrophobic cavity and one remained outside. This equilibrated structure was positioned 1nm away from the *inner* and *raft* large bilayers and simulated for 1 μ*s*. Lastly, three separate monomers with exposed Myr were positioned near the bilayer and within 2.5 nm of each other to examine their interaction with other protein units as well as with the bilayer without any bias. Overall, we simulated 13 systems for a total of 42.8 μ*s*, including the short 200ns trajectories of an MA monomer on the small membrane patches.

Simulations were carried using isobaric and isothermal dynamics (constant NPT ensemble) with a simulation timestep of 2fs and periodic boundary conditions. The temperature was kept constant at 310.15 K using the Nose-Hoover thermostat (45, 46) with a coupling time constant of 1.0 ps in GROMACS. Similarly, the pressure was set at 1bar and controlled using the Parrinello-Rahman barostat semi-isotropically with a compressibility of 4.5 × 10_-5_ and a coupling time constant of 5.0 ps (47, 48). Van Der Waals interactions were computed using a switching function between 1.0 and 1.2 nm, and long-range electrostatics evaluated using Particle Mesh Ewald (49). Hydrogen bonds in GROMACS were constrained using the LINCS algorithm (50). Simulation parameters for the Anton2 machine are set by its ark guesser files (scripts designed to optimize the parameters for the integration algorithms to enhance numerical integration during the simulation); as such, the cut-off values to compute interactions between neighboring atoms are selected automatically during system preparation. Long-range electrostatics were computed using the Gaussian Split Ewald algorithm (51), and hydrogen bonds constrained using the SHAKE algorithm (52). Finally, the Nose-Hoover thermostat and MTK barostat control the temperature and pressure respectively during NPT dynamics using optimized parameters set by the *Multigrator* integrator on Anton2 (53).

Analysis of the trajectories was performed using GROMACS, MDAnalysis (54, 55), MDTraj (56), and PyEmma (57) packages. Visual Molecular Dynamics (VMD) software package (58) was used to visualize the systems and generate all the snapshots. Description of the analysis included in the next sections is summarized on Table S4. All reported values are blocked averages with their respective standard error for each given quantity over at least 100ns of equilibrated trajectory unless stated otherwise. In general, distances were computed between the center-of-mass of the corresponding atom(s); angles were computed defining vectors between the center-of-mass of the corresponding initial and final residues; the bilayer center was estimated as the center-of-mass between the phosphorus atoms in each leaflet leaflets. Time series plots show the raw data faded in the background and the moving average, computed every 30 data points, as bolded lines for clarity. Lastly, we carried time-lagged independent component analysis (tICA) (59-61) as a reduction technique to quantify the protein changes during Myr insertion. We selected the two slowest independent components (tICs) computed as the linear combination of the distance between alpha carbons in the protein and the carbon atoms of the Myr tail. As an estimate of the relative contributions of the tICs to the Myr conformation or state, i.e. sequestered, exposed, inserted, we computed the natural log of the probability densities of the projected trajectories onto the corresponding tIC.

## V. Conclusions

All atom MD simulations were used to examine specific interactions between the MA domain of HIV-1 Gag polyprotein with lipid bilayers and to determine the effect of bilayer nature on the binding events as well as insertion of Myr, the lipidated tail of MA. We simulated an MA monomer near the surface of small membrane patches, three separate monomers on larger membrane patches, and a formed MA trimer. Our studies show MA binding to the PM is largely stabilized by electrostatic interactions between the HBR and charged lipid headgroups, specially PIP_2_ lipids. Myr insertion is not needed to anchor the protein to the binding leaflet, but it has an impact on the lateral organization of lipids in the bilayer. In our simulations we observed two bound conformations, but only one that is feasible in the case when the rest of the Gag polyprotein is present: the open conformation with the mouth of the protein pointing sideways. This bound conformation results in the HBR laying horizontally on the bilayer, the recruitment of PIP_2_ lipids to the binding site, and provides enough space for Myr release and insertion into the bilayer. The open conformation is the only one that allows Myr insertion into the binding leaflet in what seems to be its preferred mechanism, by unrolling of the Myr lipid tail into the leaflet.

Our simulations are the first of their kind to capture repeatedly the insertion of the Myr tail through unbiased all-atom MD simulations. Through our analysis we have shown the time scale for insertion is on average 50 ns, and only occurs when the protein is bound in the open conformation. As Myr unrolls into the binding leaflet, the first helix in the protein sequence, H1, shifts to rest flat on the membrane surface. The first and last carbons in the Myr tail, C2 and C14 respectively, pass each other as the tail inserts into the leaflet; depending on the location of Myr prior to insertion, C14 can extend vertically towards the solvent or enter the leaflet “sweeping” the membrane surface as Myr unrolls into the leaflet from a flat position between the protein and the membrane. This mechanism was further characterized using tICA, identifying the HBR, H1, and H5 as key protein regions that modulate or contribute to the insertion process. We simulated three membrane models to capture different membrane mechanical and structural properties, and found Myr only inserts into our *inner* membrane model, that mimics the inner leaflet of the PM in terms of anionic lipid and sterol content.

The protein was randomly positioned near the membrane surface after a short equilibration in solvent, so we have not quantified the effect of lipid domains on initial protein binding or Myr insertion. Such a study would provide additional insights on the preference of MA for lipid order or disordered domains. We have shown, however, that insertion of Myr results in modifications of lipid order as quantified by the order parameter S_CD_. The effect is not as dramatic for the case of a single monomer, which alters the order of the last carbons in the long sphingomyelin lipid tails. However, the presence of a formed trimer that enables quicker Myr insertion, induces increase in order in the mid-region of the same lipids. Our results present evidence that an increasing number of protein units enhances the changes observed on the membrane upon MA binding, as would be the case at early stages of the viral assembly process of HIV-1 in the cell.

One of the main characteristics of the PM is the asymmetry between its bilayers; here, we examined symmetric membrane models and reserved a study with asymmetric models for a separate publication. Conclusions from this present work serve to reveal the specific MA-lipid contacts, effects of membrane nature on protein binding and Myr insertion events, and membrane response to protein binding. The MA domain is a conserved domain of Gag across several retroviruses, thus we expect our conclusions to also provide insight to understand the mechanism of binding in other viral systems (62). Fundamental understanding of the membrane targeting mechanism of MA can offer new perspectives for inhibitor-based anti-retroviral treatments (63) and expand the general understanding about peripheral protein interactions with membranes and corresponding implications in macromolecular assembly.

## Supporting information

Supplemental References

## Author Contributions

V. Monje-Galvan carried all the simulations and analysis of the systems reported in this work and was in charge of manuscript preparation under the advisement and mentoring of G.A. Voth. The authors thank Drs. A. J. Pak, F. Aydin, S. Kim, and A. Yu for consultation regarding the tICA methodology and general discussions on HIV-1 dynamics throughout the duration of the study and manuscript preparation.

## Acknowledgements

This work was funded by the National Institute of General Medical Sciences of the United States NIH under grant R01GM063796. Computational resources were provided by the Pittsburgh Super Computing Center through the Anton 2 machine under Grant R01GM116961 from the National Institutes of Health, and the specific allocation PSCA17046P. The Anton 2 machine at PSC was generously made available by D.E. Shaw Research. Part of this work was also completed with resources from the University of Chicago Research Computing Center, and the Extreme Science and Engineering Discovery Environment, supported by the National Science Foundation grant number ACI-1548562.

## References

1. Khan, Hanif M., T. He, E. Fuglebakk, C. Grauffel, B. Yang, Mary F. Roberts, A. Gershenson, and N. Reuter. 2016. A Role for Weak Electrostatic Interactions in Peripheral Membrane Protein Binding. Biophysical Journal 110(6):1367–1378.

2. Whited, A. M., and A. Johs. 2015. The interactions of peripheral membrane proteins with biological membranes. Chemistry and Physics of Lipids 192:51–59.

3. Barros, M., F. Heinrich, S. A. Datta, A. Rein, I. Karageorgos, H. Nanda, and M. Losche. 2016. Membrane Binding of HIV-1 Matrix Protein: Dependence on Bilayer Composition and Protein Lipidation. J Virol 90(9):4544–4555.

4. Munro, S. 2002. Organelle identity and the targeting of peripheral membrane proteins. Current Opinion in Cell Biology 14(4):506–514.

5. Monje-Galvan, V., and J. B. Klauda. 2016. Peripheral membrane proteins: Tying the knot between experiment and computation. Biochimica et Biophysica Acta (BBA) - Biomembranes 1858(7, Part B):1584–1593.

6. Sens, P., L. Johannes, and P. Bassereau. 2008. Biophysical approaches to protein-induced membrane deformations in trafficking. Current Opinion in Cell Biology 20(4):476–482.

7. Stahelin, R. V. 2013. Chapter 19 - Monitoring Peripheral Protein Oligomerization on Biological Membranes. Methods in Cell Biology. P. M. Conn, editor. Academic Press, pp. 359–371.

8. Simunovic, M., P. Bassereau, and G. A. Voth. 2018. Organizing membrane-curving proteins: the emerging dynamical picture. Current Opinion in Structural Biology 51:99–105.

9. Rogaski, B., and J. B. Klauda. 2012. Membrane-Binding Mechanism of a Peripheral Membrane Protein through Microsecond Molecular Dynamics Simulations. Journal of Molecular Biology 423(5):847–861.

10. de la Ballina, L. R., M. J. Munson, and A. Simonsen. 2019. Lipids and Lipid-Binding Proteins in Selective Autophagy. Journal of Molecular Biology.

11. Kerr, D., G. T. Tietjen, Z. Gong, E. Tajkhorshid, E. J. Adams, and K. Y. C. Lee. 2018. Sensitivity of peripheral membrane proteins to the membrane context: A case study of phosphatidylserine and the TIM proteins. Biochimica et Biophysica Acta (BBA) - Biomembranes 1860(10):2126–2133.

12. Olety, B., and A. Ono. 2014. Roles played by acidic lipids in HIV-1 Gag membrane binding. Virus Res 193:108–115.

13. Sengupta, P., A. Y. Seo, H. A. Pasolli, Y. E. Song, M. Johnson, and J. Lippincott-Schwartz. 2019. A lipid-based partitioning mechanism for selective incorporation of proteins into membranes of HIV particles. Nature Cell Biology 21(4):452–461.

14. Huber, R. G., J. K. Marzinek, D. A. Holdbrook, and P. J. Bond. 2017. Multiscale molecular dynamics simulation approaches to the structure and dynamics of viruses. Prog Biophys Mol Biol 128:121–132.

15. Dick, R. A., and V. M. Vogt. 2014. Membrane interaction of retroviral Gag proteins. Virology 5.

16. Lalonde, M. S., and W. I. Sundquist. 2012. How HIV finds the door. Proc Natl Acad Sci U S A 109(46):18631–18632.

17. Mercredi, P. Y., N. Bucca, B. Loeliger, C. R. Gaines, M. Mehta, P. Bhargava, P. R. Tedbury, L. Charlier, N. Floquet, D. Muriaux, C. Favard, C. R. Sanders, E. O. Freed, J. Marchant, and M. F. Summers. 2016. Structural and Molecular Determinants of Membrane Binding by the HIV-1 Matrix Protein. J Mol Biol 428(8):1637–1655.

18. Vlach, J., and J. S. Saad. 2015. Structural and molecular determinants of HIV-1 Gag binding to the plasma membrane. Front Microbiol 6:232.

19. Tedbury, P. R., M. Novikova, S. D. Ablan, and E. O. Freed. 2016. Biochemical evidence of a role for matrix trimerization in HIV-1 envelope glycoprotein incorporation. Proc Natl Acad Sci U S A 113(2):E182–190.

20. Yandrapalli, N., Q. Lubart, H. S. Tanwar, C. Picart, J. Mak, D. Muriaux, and C. Favard. 2016. Self assembly of HIV-1 Gag protein on lipid membranes generates PI(4,5)P2/Cholesterol nanoclusters. Sci Rep 6:39332.

21. O’Carroll, I. P., F. Soheilian, A. Kamata, K. Nagashima, and A. Rein. 2013. Elements in HIV-1 Gag contributing to virus particle assembly. Virus Res 171(2):341–345.

22. Alfadhli, A., R. L. Barklis, and E. Barklis. 2009. HIV-1 matrix organizes as a hexamer of trimers on membranes containing phosphatidylinositol-(4,5)-bisphosphate. Virology 387(2):466–472.

23. Eells, R., M. Barros, K. M. Scott, I. Karageorgos, F. Heinrich, and M. Losche. 2017. Structural characterization of membrane-bound human immunodeficiency virus-1 Gag matrix with neutron reflectometry. Biointerphases 12(2):02D408.

24. Pak, A. J., J. M. A. Grime, P. Sengupta, A. K. Chen, A. E. P. Durumeric, A. Srivastava, M. Yeager, J. A. G. Briggs, J. Lippincott-Schwartz, and G. A. Voth. 2017. Immature HIV-1 lattice assembly dynamics are regulated by scaffolding from nucleic acid and the plasma membrane. Proc Natl Acad Sci U S A 114(47):E10056–E10065.

25. Leung, K., J.-O. Kim, L. Ganesh, J. Kabat, O. Schwartz, and G. J. Nabel. 2008. HIV-1 Assembly: Viral Glycoproteins Segregate Quantally to Lipid Rafts that Associate Individually with HIV-1 Capsids and Virions. Cell Host & Microbe 3(5):285–292.

26. Metzner, C., B. Salmons, W. H. Günzburg, and J. A. Dangerfield. 2008. Rafts, anchors and viruses — A role for glycosylphosphatidylinositol anchored proteins in the modification of enveloped viruses and viral vectors. Virology 382(2):125–131.

27. Lorent, J. H., and I. Levental. 2015. Structural determinants of protein partitioning into ordered membrane domains and lipid rafts. Chem Phys Lipids 192:23–32.

28. Saliba, A. E., I. Vonkova, and A. C. Gavin. 2015. The systematic analysis of protein-lipid interactions comes of age. Nat Rev Mol Cell Biol 16(12):753–761.

29. Monje-Galvan, V., and J. B. Klauda. 2015. Modelling Yeast Organelle Membranes and How Lipid Diversity influences Bilayer Properties. Biochemistry 54:6852–6861.

30. Bukrinskaya, A. 2007. HIV-1 matrix protein: A mysterious regulator of the viral life cycle. Virus Research 124(1):1–11.

31. Alfadhli, A., and E. Barklis. 2014. The roles of lipids and nucleic acids in HIV-1 assembly. Front Microbiol 5:253.

32. Saad, J. S., J. Miller, J. Tai, A. Kim, R. H. Ghanam, and M. F. Summers. 2006. Structural basis for targeting HIV-1 Gag proteins to the plasma membrane for virus assembly. Proceedings of the National Academy of Sciences 103(30):11364.

33. Charlier, L., M. Louet, L. Chaloin, P. Fuchs, J. Martinez, D. Muriaux, C. Favard, and N. Floquet. 2014. Coarse-grained simulations of the HIV-1 matrix protein anchoring: revisiting its assembly on membrane domains. Biophys J 106(3):577–585.

34. Tang, C., E. Loeliger, P. Luncsford, I. Kinde, D. Beckett, and M. F. Summers. 2004. Entropic switch regulates myristate exposure in teh HIV-1 matrix protein. Proc Natl Acad Sci U S A 101(2):517–522.

35. Jo, S., T. Kim, and W. Im. 2007. Automated builder and database of protein/membrane complexes for molecular dynamics simulations. PLoS One 2(9):e880.

36. Jo, S., T. Kim, V. G. Iyer, and W. Im. 2008. CHARMM-GUI: A web-based graphical user interface for CHARMM. Journal of Computational Chemistry 29(11):1859–1865.

37. Brooks, B. R., C. L. Brooks, 3rd, A. D. Mackerell, Jr., L. Nilsson, R. J. Petrella, B. Roux, Y. Won, G. Archontis, C. Bartels, S. Boresch, A. Caflisch, L. Caves, Q. Cui, A. R. Dinner, M. Feig, S. Fischer, J. Gao, M. Hodoscek, W. Im, K. Kuczera, T. Lazaridis, J. Ma, V. Ovchinnikov, E. Paci, R. W. Pastor, C. B. Post, J. Z. Pu, M. Schaefer, B. Tidor, R. M. Venable, H. L. Woodcock, X. Wu, W. Yang, D. M. York, and M. Karplus. 2009. CHARMM: the biomolecular simulation program. J Comput Chem 30(10):1545–1614.

38. Wu, E. L., X. Cheng, S. Jo, H. Rui, K. C. Song, E. M. Davila-Contreras, Y. Qi, J. Lee, V. Monje-Galvan, R. M. Venable, J. B. Klauda, and W. Im. 2014. CHARMM-GUI Membrane Builder toward realistic biological membrane simulations. Journal of Computational Chemistry.

39. Lee, J., X. Cheng, J. M. Swails, M. S. Yeom, P. K. Eastman, J. A. Lemkul, S. Wei, J. Buckner, J. C. Jeong, Y. Qi, S. Jo, V. S. Pande, D. A. Case, C. L. Brooks, A. D. MacKerell, J. B. Klauda, and W. Im. 2016. CHARMM-GUI Input Generator for NAMD, GROMACS, AMBER, OpenMM, and CHARMM/OpenMM Simulations Using the CHARMM36 Additive Force Field. Journal of Chemical Theory and Computation 12(1):405–413.

40. Huang, J., S. Rauscher, G. Nawrocki, T. Ran, M. Feig, B. L. de Groot, H. Grubmüller, and A. D. MacKerell. 2017. CHARMM36m: an improved force field for folded and intrinsically disordered proteins. Nature Methods 14(1):71–73.

41. Abraham, M. J., T. Murtola, R. Schulz, S. Páll, J. C. Smith, B. Hess, and E. Lindahl. 2015. GROMACS: High performance molecular simulations through multi-level parallelism from laptops to supercomputers. SoftwareX 1-2:19–25.

42. Shaw, D. E., J. P. Grossman, J. A. Bank, B. Batson, J. A. Butts, J. C. Chao, M. M. Deneroff, R. O. Dror, A. Even, C. H. Fenton, A. Forte, J. Gagliardo, G. Gill, B. Greskamp, C. R. Ho, D. J. Ierardi, L. Iserovich, J. S. Kuskin, R. H. Larson, T. Layman, L. Lee, A. K. Lerer, C. Li, D. Killebrew, K. M. Mackenzie, S. Y. Mok, M. A. Moraes, R. Mueller, L. J. Nociolo, J. L. Peticolas, T. Quan, D. Ramot, J. K. Salmon, D. P. Scarpazza, U. B. Schafer, N. Siddique, C. W. Snyder, J. Spengler, P. T. P. Tang, M. Theobald, H. Toma, B. Towles, B. Vitale, S. C. Wang, and C. Young. 2014. Anton 2: Raising the Bar for Performance and Programmability in a Special–Purpose Molecular Dynamics Supercomputer. In SC ‘14: Proceedings of the International Conference for High Performance Computing, Networking, Storage and Analysis. 41–53.

43. Jo, S., J. B. Lim, J. B. Klauda, and W. Im. 2009. CHARMM-GUI Membrane Builder for mixed bilayers and its application to yeast membranes. Biophys J 97(1):50–58.

44. Hill, C. P., D. Worthylake, D. P. Bancroft, A. M. Christensen, and W. I. Sundquist. 1996. Crystal structures of the trimeric human immonodeficiency virus type 1 matrix protein: Implications for membrane association and assembly. Proc Natl Acad Sci U S A 93:3099–3104.

45. Hoover, W. G. 1985. Canonical dynamics: Equilibrium phase-space distributions. Phys Rev A Gen Phys 31(3):1695–1697.

46. Nosé, S. 1984. A molecular dynamics method for simulations in the canonical ensemble. Mol Phys 52:255–268.

47. Nosé, S., and M. L. Klein. 1983. Constant pressure molecular dynamics for molecular systems. Mol Phys 50:1055–1076.

48. Parrinello, M., and A. Rahman. 1981. Polymorphic transitions in single crystals: A new molecular dynamics method. J Appl Phys 52:7182–7190.

49. Darden, T., D. York, and L. Pedersen. 1993. Particle Mesh Ewald - an N.Log(N) Method for Ewald Sums in Large Systems. Journal of Chemical Physics 98(12):10089–10092.

50. Hess, B., H. Bekker, H. J. C. Berendsen, and J. G. E. M. Fraaije. 1997. LINCS: A linear constraint solver for molecular simulations. J Comput Chem 18:1463–1472.

51. Shan, Y., J. L. Klepeis, M. P. Eastwood, R. O. Dror, and D. E. Shaw. 2005. Gaussian split Ewald: A fast Ewald mesh method for molecular simulation. J Chem Phys 122(5):54101.

52. Ryckaert, J.-P., G. Ciccotti, and H. J. C. Berendsen. 1977. Numerical integration of the cartesian equations of motion of a system with constraints: molecular dynamics of n-alkanes. Journal of Computational Physics 23(3):327–341.

53. Lippert, R. A., C. Predescu, D. J. Ierardi, K. M. Mackenzie, M. P. Eastwood, R. O. Dror, and D. E. Shaw. 2013. Accurate and efficient integration for molecular dynamics simulations at constant temperature and pressure. J Chem Phys 139(16):164106. Text.

54. Gowers, R. J., M. Linke, J. Barnoud, T. J. Reddy, M. N. Melo, S. L. Seyler, J. D. Domański, D. L. Dotson, S. Buchoux, I. M. Kenney, and O. Beckstein. 2016. MDAnalysis: A Python Package for the Rapida Analysis of Molecualr Dynamics Simulations. Proceedings of the 15th Python in Science Conference:98–105.

55. Michaud-Agrawal, N., E. J. Denning, T. B. Woolf, and O. Beckstein. 2011. MDAnalysis: A toolkit for the analysis of molecular dynamics simulations. Journal of Computational Chemistry 32(10):2319–2327.

56. McGibbon, Robert T., Kyle A. Beauchamp, Matthew P. Harrigan, C. Klein, Jason M. Swails, Carlos X. Hernández, Christian R. Schwantes, L.-P. Wang, Thomas J. Lane, and Vijay S. Pande. 2015. MDTraj: A Modern Open Library for the Analysis of Molecular Dynamics Trajectories. Biophysical Journal 109(8):1528–1532.

57. Scherer, M. K., B. Trendelkamp-Schroer, F. Paul, G. Pérez-Hernández, M. Hoffmann, N. Plattner, C. Wehmeyer, J.-H. Prinz, and F. Noé. 2015. PyEMMA 2: A Software Package for Estimation, Validation, and Analysis of Markov Models. Journal of Chemical Theory and Computation 11(11):5525–5542.

58. Humphrey, W., A. Dalke, and K. Schulten. 1996. VMD: visual molecular dynamics. J Mol Graph 14(1):33-38, 27-38.

59. Pérez-Hernández, G., and F. Noé. 2016. Hierarchical Time-Lagged Independent Component Analysis: Computing Slow Modes and Reaction Coordinates for Large Molecular Systems. Journal of Chemical Theory and Computation 12(12):6118–6129.

60. Pérez-Hernández, G., F. Paul, T. Giorgino, G. De Fabritiis, and F. Noé. 2013. Identification of slow molecular order parameters for Markov model construction. The Journal of Chemical Physics 139(1):015102.

61. M. Sultan, M., and V. S. Pande. 2017. tICA-Metadynamics: Accelerating Metadynamics by Using Kinetically Selected Collective Variables. Journal of Chemical Theory and Computation 13(6):2440–2447.

62. Mattei, S., A. Tan, B. Glass, B. Müller, H.-G. Kräusslich, and J. A. G. Briggs. 2018. High-resolution structures of HIV-1 Gag cleavage mutants determine structural switch for virus maturation. Proceedings of the National Academy of Sciences 115(40):E9401.

63. Zentner, I., L. J. Sierra, L. Maciunas, A. Vinnik, P. Fedichev, M. K. Mankowski, R. G. Ptak, J. Martin-Garcia, and S. Cocklin. 2013. Discovery of a small-molecule antiviral targeting the HIV-1 matrix protein. Bioorg Med Chem Lett 23(4):1132–1135.

