## Supplemental Table S1 for "Binding Mechanism of the Matrix Domain of HIV-1 Gag on Lipid Membranes"

**Table S1.** Simulation components for the membrane models used in this study. Chemical structures of each lipid species are shown in Fig S1.

| **Model** | **Lipid species** | **Lipid mol fract.** | **# waters (small)** | **Ions** | **# waters (large)** | **Ions** |
|---|---|---|---|---|---|---|
| Chol-free<br>*APL* 66.9 +/- 0.84* | DOPC<br>DOPS<br>$PIP_2$+ | 0.80<br>0.15<br>0.05 | 23442<br>*78.1 H-num*** | 211 K<br>112 Cl | 23442<br>*78.1 H-num*** | 211 K<br>112 Cl |
| Inner<br>*APL 47.8 +/- 0.40* | Chol<br>DOPC<br>DOPS<br>POPE<br>BSM<br>$PIP_2$ | 0.30<br>0.17<br>0.17<br>0.25<br>0.08<br>0.02 | 16009<br>*53.4 H-num* | 154 K<br>81 Cl | 16009<br>*53.4 H-num* | 154 K<br>81 Cl |
| Raft<br>*APL 47.1 +/- 0.40* | Chol<br>DOPC<br>DOPS<br>BSM<br>$PIP_2$ | 0.28<br>0.30<br>0.06<br>0.30<br>0.06 | 15744<br>*52.6 H-num* | 170 K<br>83 Cl | 15744<br>*52.6 H-num* | 170 K<br>83 Cl |

**area per lipid.* ***hydration number (# waters/lipid).* +*SAPI25 (20:4 – 18:0)*
