## Supplemental Table S2 for "Binding Mechanism of the Matrix Domain of HIV-1 Gag on Lipid Membranes"

**Table S2.** Matrix bound times and conformation for the 200ns equilibration trajectories

| System | Bound leaflet | Conformation | 1st contact (ns) | Myr release (ns) |
|---|---|---|---|---|
| *Chol-free*$_{200}$ | top | open | 90 ± 1 | 112 ± 2 |
| *Chol-free-1*$_{200}$ | bottom | open | 68 ± 1 | 80 ± 2 |
| *Inner*$_{200}$ | bottom | open | 52 ± 1 | 145 ± 2 |
| *Inner-1*$_{200}$ | top | blocked | 70 ± 1 | *no release* |
| *Raft*$_{200}$ | top | blocked | 25 ± 1 | *no release* |
| *Raft-1*$_{200}$ | top | open | 35 ± 1 | 75 ± 2 |
