## Supplemental Table S3 for "Binding Mechanism of the Matrix Domain of HIV-1 Gag on Lipid Membranes"

**Table S3.** Simulation details for the microsecond trajectories of MA proteins

| ***Monomers*** | | | | |
|---|---|---|---|---|
| **System** | **length (ns)** | **Bound leaflet** | **Conformation** | **Myr location** |
| *Chol-free$_{pre}$* | 5000 | top | open | *pre-inserted* |
| *Chol-free-1$_{pre}$* | 3000 | top | open | *pre-inserted* |
| *Chol-free-2$_{pre}$* | 5000 | top | | *pre-inserted* |
| *Inner* | 5000 | bottom | open | exposed |
| *Inner-1* | 1600 | bottom | open | exposed |
| *Inner$_{myr}$* | 5000 | top | open | partially-inserted |
| *Raft* | 5000 | top | blocked | hydrophobic cavity |
| *Raft-1* | 3000 | top | blocked | hydrophobic cavity |
| *Raft$_{pre}$* | 5000 | top | open/blocked | *pre-inserted* |
| ***Three units*** | | | | |
| **System** | **length (ns)** | **Bound leaflet** | **Conformation** | **Myr insertion** |
| *inner-L$_{trimer}$* | 1000 | top | trimer | yes (2) |
| *inner-L$_{mono}$* | 1000 | top | 3 monomers | none |
| *raft-L$_{trimer}$* | 1000 | top | trimer | none |
| *raft-L$_{mono}$* | 1000 | top | 3 monomers | none |
