## Supplemental Table S4 for "Binding Mechanism of the Matrix Domain of HIV-1 Gag on Lipid Membranes"

**Table S4.** Analyses performed on the systems to quantify protein binding, Myr insertion, protein conformational changes, membrane mechanical/structural properties. (The trajectories were re-centered around the protein and the binding leaflet using GROMACS(1) *trajectorv* tool prior to computing the respective quantities.)

| Analysis | Description | Systems analyzed |
|---|---|---|
| **Distances** (Å)<br>*MDAnalysis(2, 3)* | In most of the cases the z-component of the distance between the center-of-mass (com) of molecules was computed. The bilayer center was determined as the average position between the phosphorus atoms of each leaflet. For the Lid-H1 distance, the absolute value of the distance is reported. | *All systems* |
| **Self-distance** (Å)<br>*MDAnalysis*<br>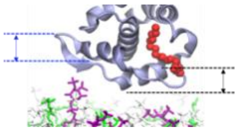 | We defined this quantity as the z-component of the difference between the position of the last and first alpha carbons of a sequence of residues in the protein. For H1, the difference was computed between the com of residues 18 and 8 as shown with the black bashed lines in the sketch to the left.<br>For the HBR, the middle residue (25) in the sequence was taken as the "last" residue for this calculation since this region is a U-loop; the first residue was defined as the com between the actual first and last residues in the sequence (19 & 29). This distance is shown with the blue dashed lines in the sketch.<br>A positive value indicates the last residue in the sequence is higher than the first; a value of zero indicates the section is essentially flat. | *Inner, inner1, inner-L$_{trimer}$* |
| **Cosine of the angle**<br>*MDAnalysis* | The absolute value of the cosine of the angle between a vector defined for the Myr tail and the unit vector in the z-direction is computed and reported. We took the absolute value to present a unified measurement of the orientation of Myr in the bilayer given the protein indistinctly binds the top or bottom leaflets of our symmetric models. The Myr vector is defined from the com of C2 to the com of C14, the first and last atoms of Myr.<br>A value of 1 indicates Myr is aligned with the z-axis, parallel to the bilayer normal; a value of 0 indicates it is perpendicular to it, i.e. is aligned to the membrane surface. | *Inner, inner1* and *inner-L$_{trimer}$* systems; all *chol-free$_{pre}$* and *raft$_{pre}$* systems. The *inner$_{myr}$* replica is also reported as reference. |
| **Lipid counts**<br>*MDAnalysis* | Count of the number of phosphorus atoms from $PIP_2$ and DOSP lipid species within 1 nm of the alpha carbons of the protein. | *All systems* |
| **Lipid densities**<br>*MDTraj (4)* | 2D histograms of the x-y position of each phosphorus atom in the lipid species of interest computed during the last 50ns of simulation for the short trajectories, and the last 300 ns for the microsecond trajectories. | *All systems* |
| **S$_{CD}$ parameters**<br>*VMD (5)* | The lipid tail order parameters serve to characterize the structure and order of the lipid tails inside the bilayer. They are computed from simulation data measuring the angle $\theta$ between every C-H bond in the lipid tails and the bilayer normal (z-axis) according to:<br>$$S_{CD} = \left|{}^{3}/_{2}\, cos^2\theta - {}^{1}/_{2}\right|$$<br>Higher values indicate a more ordered bilayer; lower values are expected for the last carbons in the lipid tails and regions with double bonds. Increased sterol concentration tends to increase order in the bilayer (6). The order parameters were computed over the last 200ns of the equilibrated trajectory. | *Inner, inner1, inner-L$_{trimer}$* |

| | | |
|---|---|---|
| **Area per lipid** (Å2/lipid) *GROMACS* | The average surface area of the x-y plane (parallel to the membrane surface) was computed from the simulation box dimensions over the last 50ns of simulation of the membrane-only equilibration runs of the small membrane patches. The value was divided among the lipids in each leaflet (150) to obtain the average APL for each model and is reported on Table S1. | *Chol-free*, *inner*, *raft* (small membrane patches) |
| **RMSF** *MDAnalysis* | The root-mean-square-fluctuations (RMSF) were computed per residue using the *RMSF* tool from *MDAnalysis* to measure the protein changes during Myr insertion. During the analysis, the protein is aligned to the its structure in the first frame of the trajectory to avoid including translation and rotation of the protein in the calculation. | *All systems* |
| **tICA** *VMD, MDtraj, MDAnalysis, PyEmma (7)* | Time-lagged independent component analysis is a reduction of dimensionality technique to obtain the slowest independent component in a given process as linear combinations of *n* features. These components are identified by solving the eigenvalue-eigenvector problem for the co-variance matrix of the desired process (8-11). To characterize the Myr insertion in our simulations we examined the distances between every carbon in the Myr tail and the $C_\alpha$ in the main regions of the protein, i.e. H1-H5 and the HBR (all the other coiled regions were not considered). This feature was selected as the best description for Myr insertion after comparison with other features such as the cartesian coordinates of the protein, distances among Myr carbon atoms, and the protein RMSD. The trajectory was pre-processed and the protein aligned to its position in the first frame of the trajectory using VMD to prevent translational and rotational degrees of freedom from interfering in tICA. The lag-time for this analysis was set to 50 after analyzing the decay of eigenvalues with respect to lag time for the two slowest tICs. The coefficients of the corresponding eigenvectors, a.k.a. tIC loads, were also examined to determine the distances that contribute more to the respective tIC, i.e. the slowest motions during Myr insertion. | *Inner*, *Inner-1*, *inner-L* w/trimer systems, and the small *chol-free* and *raft* systems with no Myr insertion. |
| **tIC contributions to Myr conformation** (energy/kT) *MDAnalysis, PyEmma, Seaborn (12)* | An estimate of the relative contributions of a given metric to the state or conformation of Myr in the system, we computed the logarithm of the histogrammed data of (i) the C2 and C14 distances from the bilayer center, see Fig. 4 in the main manuscript; and (ii) the slowest independent components identified from *tICA.* The estimates are given in units of kT (7), a factor of 2.6 for T=310K would render the units in KJ/mol. To facilitate the comparison among computed values from different trajectories we set the minimum to zero. | *All systems* |
