## Supplemental References for "Binding Mechanism of the Matrix Domain of HIV-1 Gag on Lipid Membranes"

**References for the Supplemental Information**


1. Abraham, M. J., T. Murtola, R. Schulz, S. Páll, J. C. Smith, B. Hess, and E. Lindahl. 2015. GROMACS: High performance molecular simulations through multi-level parallelism from laptops to supercomputers. SoftwareX 1-2:19-25.
2. Gowers, R. J., M. Linke, J. Barnoud, T. J. Reddy, M. N. Melo, S. L. Seyler, J. D. Domański, D. L. Dotson, S. Buchoux, I. M. Kenney, and O. Beckstein. 2016. MDAnalysis: A Python Package for the Rapida Analysis of Molecualr Dynamics Simulations. Proceedings of the 15th Python in Science Conference:98-105.
3. Michaud-Agrawal, N., E. J. Denning, T. B. Woolf, and O. Beckstein. 2011. MDAnalysis: A toolkit for the analysis of molecular dynamics simulations. Journal of Computational Chemistry 32(10):2319-2327.
4. McGibbon, Robert T., Kyle A. Beauchamp, Matthew P. Harrigan, C. Klein, Jason M. Swails, Carlos X. Hernández, Christian R. Schwantes, L.-P. Wang, Thomas J. Lane, and Vijay S. Pande. 2015. MDTraj: A Modern Open Library for the Analysis of Molecular Dynamics Trajectories. Biophysical Journal 109(8):1528-1532.
5. Humphrey, W., A. Dalke, and K. Schulten. 1996. VMD: visual molecular dynamics. J Mol Graph 14(1):33-38, 27-38.
6. Monje-Galvan, V., and J. B. Klauda. 2015. Modelling Yeast Organelle Membranes and How Lipid Diversity influences Bilayer Properties. Biochemistry 54:6852-6861.
7. Scherer, M. K., B. Trendelkamp-Schroer, F. Paul, G. Pérez-Hernández, M. Hoffmann, N. Plattner, C. Wehmeyer, J.-H. Prinz, and F. Noé. 2015. PyEMMA 2: A Software Package for Estimation, Validation, and Analysis of Markov Models. Journal of Chemical Theory and Computation 11(11):5525-5542.
8. Brotzakis, Z. F., and M. Parrinello. 2019. Enhanced Sampling of Protein Conformational Transitions via Dynamically Optimized Collective Variables. Journal of Chemical Theory and Computation 15(2):1393-1398.
9. Pérez-Hernández, G., and F. Noé. 2016. Hierarchical Time-Lagged Independent Component Analysis: Computing Slow Modes and Reaction Coordinates for Large Molecular Systems. Journal of Chemical Theory and Computation 12(12):6118-6129.
10. Pérez-Hernández, G., F. Paul, T. Giorgino, G. De Fabritiis, and F. Noé. 2013. Identification of slow molecular order parameters for Markov model construction. The Journal of Chemical Physics 139(1):015102.
11. M. Sultan, M., and V. S. Pande. 2017. tICA-Metadynamics: Accelerating Metadynamics by Using Kinetically Selected Collective Variables. Journal of Chemical Theory and Computation 13(6):2440-2447.
12. Wakson, M., O. Botvinnik, D. O'Kane, P. Hobson, J. Ostblom, S. Lukauskas, D. C. Gemperline, T. Augspurger, Y. Halchenko, J. B. Cole, J. Warmenhoven, J. d. Ruiter, C. Pye, S. Hoyer, J. Vanderplas, S. Villalba, G. Kunter, E. Quintero, P. Bachant, M. Martin, K. Meyer, A. Miles, Y. Ram, T. Brunner, T. Yarkoni, M. Lee Williams, C. Evans, C. Fitzgerald, Brian, and A. Qalieh. 2018. mwaskom/seaborn: v0.9.0 (July 2018). Zenodo.
