## Supplemental Figure S1 for "Binding Mechanism of the Matrix Domain of HIV-1 Gag on Lipid Membranes"

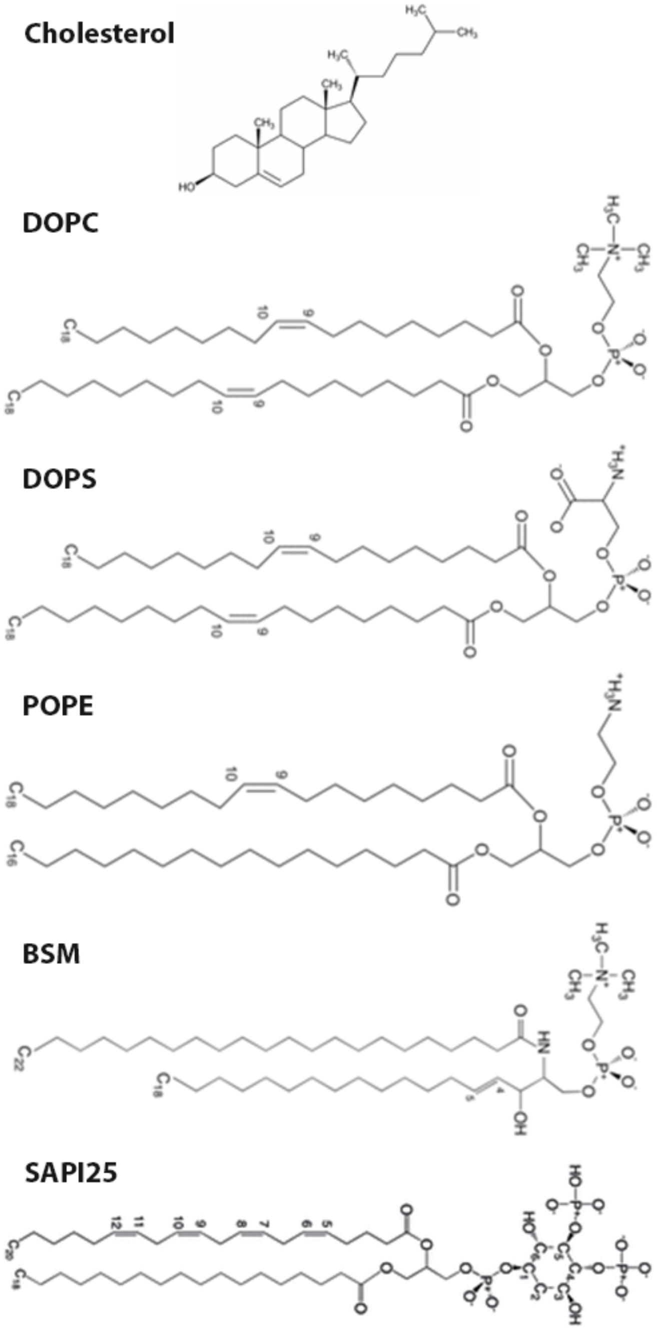


**Fig S1.** Chemical structures of the lipids used in the membrane models for this study. The lipid tail saturation nomenclature is given for the *sn2 – sn1* tails below each name.
