## Supplemental Figure S2 for "Binding Mechanism of the Matrix Domain of HIV-1 Gag on Lipid Membranes"

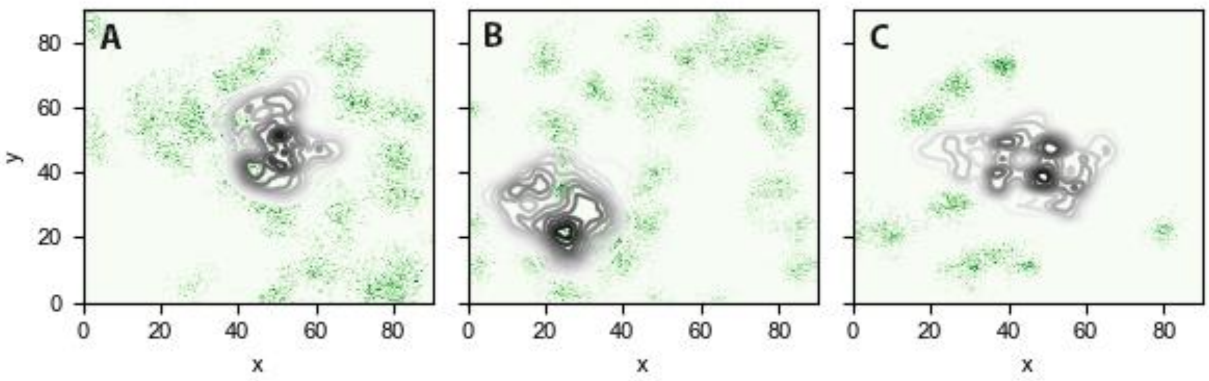


**Fig S2**. Lipid densities of DOPS lipids in the binding leaflets of the symmetric membrane models at the end of 200ns equilibration. (A) *chol-free*$_{200}$, (B) *raft*$_{200}$, and (C) *inner*$_{200}$. Densities computed over last 50ns of trajectory.
