## Supplemental Figure S3 for "Binding Mechanism of the Matrix Domain of HIV-1 Gag on Lipid Membranes"

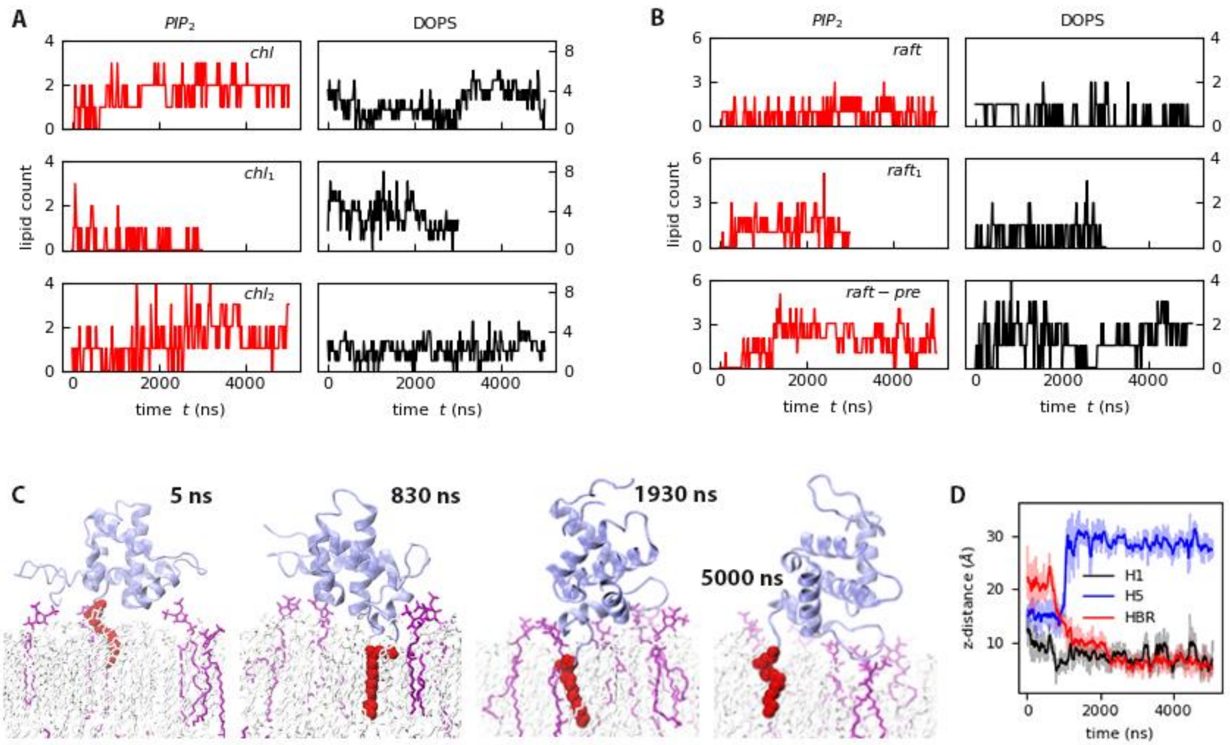


**Fig S3.** Protein-lipid contacts for the (A) *Chol-free* models with pre-inserted Myr, and (B) *Raft*, *Raft-1 and* $Raft_{pre}$ trajectories. (C) Snapshots of the protein-membrane interactions for the $raft_{pre}$ model trajectory (pre-inserted Myr), and (D) the corresponding timeseries for the distance between H1, H5, and the HBR and the bilayer center.
