## Supplemental Figure S4 for "Binding Mechanism of the Matrix Domain of HIV-1 Gag on Lipid Membranes"

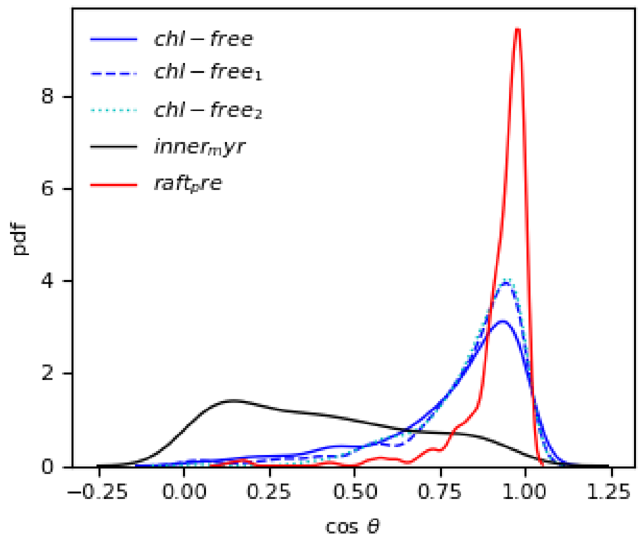


**Fig S4**. Probability distribution of the cosine of the angle between Myr and the bilayer normal (z-axis) in the systems starting from a pre-inserted Myr configuration. The $inner_{myr}$ system is also included as reference for a lipid tail that returned to the protein hydrophobic cavity and is essentially parallel to the membrane surface (black curve)
