## Supplemental Figure S5 for "Binding Mechanism of the Matrix Domain of HIV-1 Gag on Lipid Membranes"

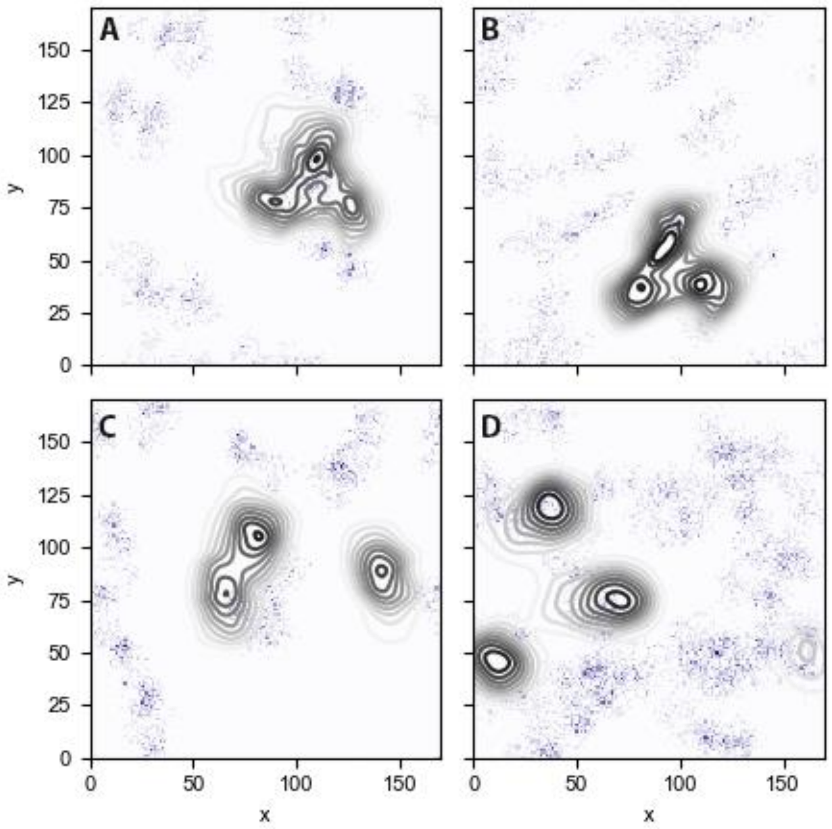


**Fig S5.** $PIP_2$ lipid densities in the binding leaflet for the: (A) *inner-$L_{trimer}$* (B) *raft-$L_{trimer}$*, (C) *inner-$L_{mono}$* (D) *raft-$L_{mono}$* systems. Data was computed over the last 200 ns of simulation.
