## Supplemental Figure S6 for "Binding Mechanism of the Matrix Domain of HIV-1 Gag on Lipid Membranes"

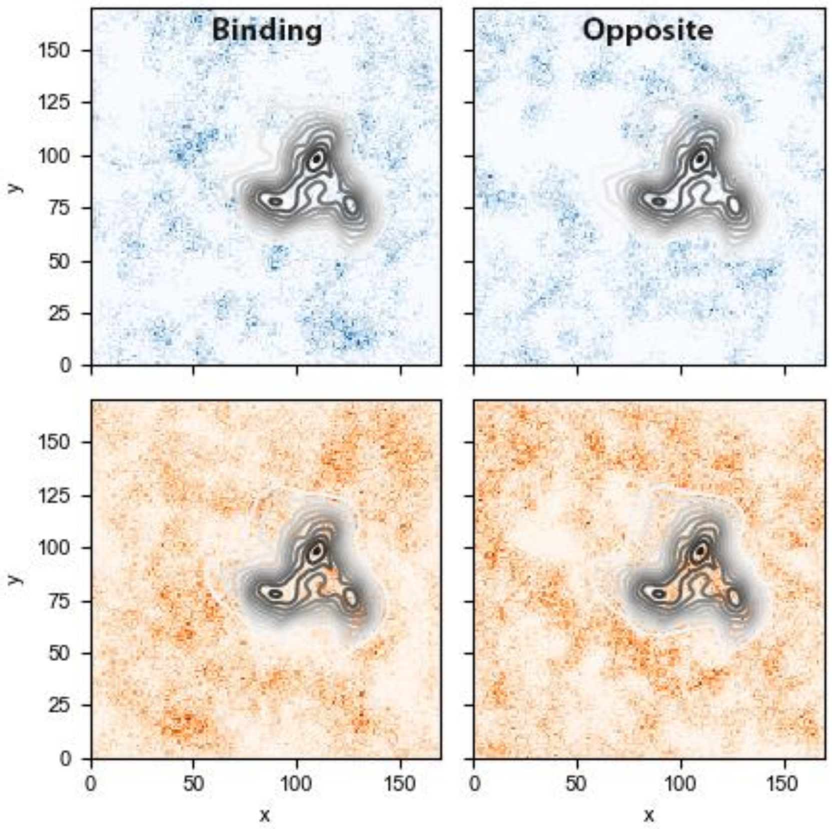


**Fig S6.** Lipid densities for sphingomyelin (top) and cholesterol (bottom) lipids in the binding and opposite leaflets, for the *inner-L*$_{trimer}$ trajectory. Data collected over the last 200 ns of simulation.
