## Supplemental Figure S7 for "Binding Mechanism of the Matrix Domain of HIV-1 Gag on Lipid Membranes"

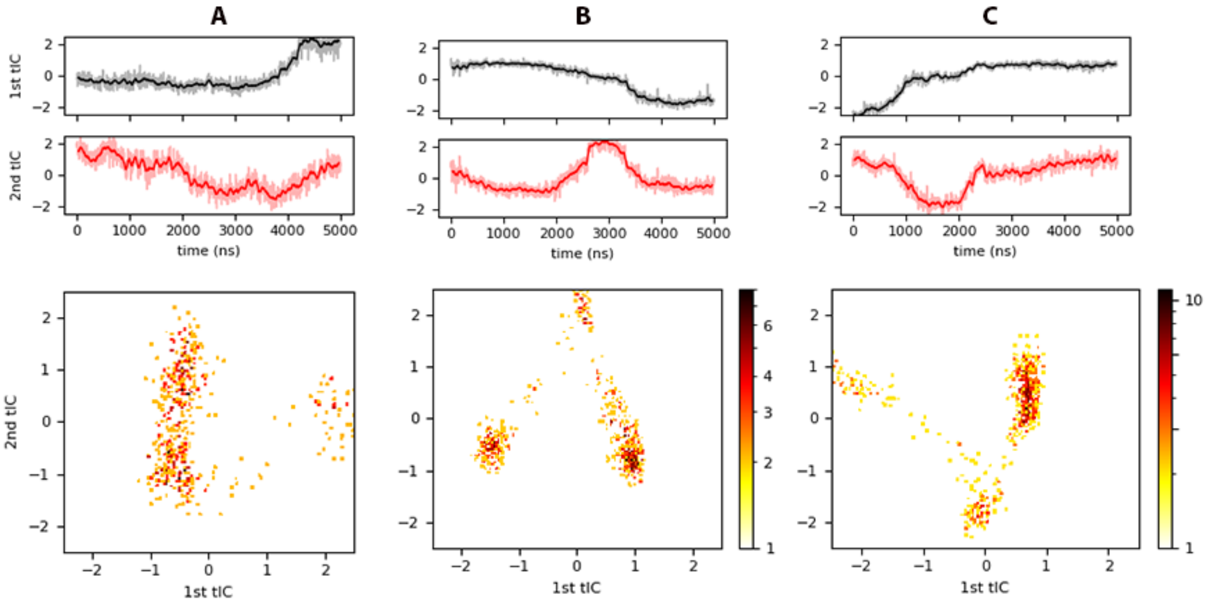


**Fig S7.** tICA results for (A) *Chol-free$_{pre}$*, (B) *Raft*, and (C) *Raft$_{pre}$* trajectories. Top panels show the projection of each trajectory onto the slowest independent components identified, and the bottom panels are heat-maps in logarithmic scale estimated from histograms of the projected trajectory onto the two slowest tICs.
